# Toward the Clinical Translation of Safe Intravenous Long Circulating Iodinated Lipid Nanoemulsion Contrast Agents for CT Imaging

**DOI:** 10.1101/2024.08.28.610138

**Authors:** Mohamed F. Attia, Ryan N. Marasco, Samuel Kwain, Charity Foxx, Daniel C. Whitehead, Alexander Kabanov, Yueh Z. Lee

**Affiliations:** Center for Nanotechnology in Drug Delivery and Division of Pharmacoengineering and Molecular Pharmaceutics, Eshelman School of Pharmacy, University of North Carolina at Chapel Hill, NC 27599, USA; Department of Chemistry, Clemson University, Clemson, SC, 29634, USA; School of Medicine, The University of North Carolina at Chapel Hill, Chapel Hill, North Carolina 27599, USA

**Author notes:** (Y. L.).

## Abstract

Current clinical small molecule x-ray CT agents are effective but pose risks such as nephrotoxicity, short blood circulation time, limiting scan durations, potential thyroid impact, and immune responses. These challenges drive the development of kidney-safe x-ray nanoparticle (NP)-based contrast agents (CAs), though translation to clinical practice is hindered by chemical complexities and potential toxicity. We have engineered an intravenous, injectable, and safe blood pool NP-based CT CAs at a clinical-equivalent dose of ∼300 mgI/kg (∼2 mL/kg), ideal for vascular and hepatic imaging which are limited by clinical agents. Our iodinated lipid nanodroplet emulsions (ILNEs) contrast agent offers high x-ray attenuation thus improved contrast enhancement, extended stability, and exceptional batch-to-batch consistency. It also boasts a straightforward and scalable manufacturing process with minimal protein interaction, prolonged blood residency (∼4h), and hepatic clearance within 3 days, avoiding nephrotoxicity. Studies in vitro, in mice, and 16.6kg porcine animal model studies confirm its safety, cytocompatibility, and absence of tissue damage. Blood, and thyroid-stimulating hormone (TSH) analyses, and kidney and liver function tests, also support further toxicity evaluations for clinical translation.

---

Advancements in medical imaging technologies, particularly Computed Tomography (CT), have revolutionized diagnostic practices by providing high-resolution, three-dimensional images crucial for detecting and monitoring diseases. Central to CT imaging are contrast agents, which enhance tissue and organ visualization. High K-edge energy contrast agents like gold nanoparticles (NPs), gadolinium, and bismuth are explored, but face challenges such as low solubility and safety concerns [1-6]. Their synthesis often involves complex, hazardous processes [7, 8].

Iodinated compounds, known for their high x-ray attenuation and biocompatibility, have been essential in clinical practice since 1950, with about 75 million doses administered annually [9]. However, traditional iodinated contrast agents have limitations, including rapid clearance, narrow imaging windows, and potential nephrotoxicity, especially in patients with compromised renal function. They can also cause adverse reactions, such as worsened renal impairment and increased risk of post-CT acute kidney injury [10-12] as well as adverse cardiac events [13, 14].

NP-based iodinated contrast agents (ICAs) may enable more targeted imaging and expand diagnostic options, but face hurdles like high excipient amounts, biological and shelf-life stability, large volume doses, production scale-up, and complex chemistry for conjugating high concentrations of iodine. Consequently, none of the developed NPs have successfully translated into clinical applications. Innovative contrast agents are needed to overcome these challenges while maintaining diagnostic efficacy and patient safety.

Lipid nanoemulsions (LNEs) have emerged as promising candidates for CT contrast enhancement [15, 16]. LNEs, characterized by nanoscale droplets of oil or lipid stabilized in an aqueous medium, offer advantages such as improved biocompatibility, tunable physicochemical properties, prolonged circulation times, reduced renal clearance, and potential for targeted delivery. LNEs mimic particles inherent in the body, like intracellular lipid droplets and low-density lipoproteins (LDLs), making them safe platforms for developing imaging agents and nanomedicines [17-19].

Incorporating iodinated compounds into LNEs can enhance x-ray attenuation, improving imaging quality while minimizing adverse effects associated with free iodinated molecules. This class of LNEs has shown high integrity in vivo and significant passive accumulation in tumor tissues via the enhanced permeability and retention (EPR) effect [20]. The key advantage is the facile surface functionalization of LNEs with ligand models or antibodies, facilitating selective targeting of specific tissues with low off-target bindings [21-24].

Translating injectable, safe, long-circulating ILNE contrast agents from preclinical studies to clinical application represents a significant milestone in molecular imaging. This transition requires thorough characterization of ILNEs’ pharmacokinetics, biodistribution, and safety profiles, along with optimizing formulation parameters to ensure efficacy and regulatory compliance.

The focus of this study is to advance the translation of new ILNE contrast agents for clinical CT imaging. By harnessing ILNEs’ unique characteristics and incorporating iodinated moieties, we aim to develop contrast agents that enhance imaging quality and exhibit improved safety profiles and pharmacokinetic properties compared to conventional counterparts. Our objectives include evaluating the in vitro stability, cytotoxicity, and hemocompatibility of the formulated contrast agents, assessing their in vivo pharmacokinetics, biodistribution, and imaging efficacy in preclinical models, and investigating safety and contrast enhancement profiles in relevant clinical settings. By fulfilling these objectives, we seek to bridge the gap between preclinical development and clinical translation of novel ILNE contrast agents, potentially enhancing diagnostic accuracy and improving patient outcomes in CT imaging.

## Materials and methods

### Materials

Non-ionic amphiphilic PEGylated surfactants: Kolliphor^®^ ELP (Cremophor^®^ EL (CrEl), a polyethoxylated (35) castor oil) and Kolliphor^®^ HS 15 (Solutol^®^ HS15, a polyethylene glycol (15)-hydroxystearate) were gifts from BASF (New York, USA). Oleic acid, 2,4,6-triiodophenol (TIPh), 4-dimethylaminopyridine (DMAP), *N,N′*-dicyclohexylcarbodiimide (DCC), dichloromethane, ethyl acetate, cyclohexane, sodium hydrogen carbonate (NaHCO_3_), sodium sulfate anhydrous (Na_2_SO_4_), sodium chloride (NaCl), deuterated chloroform (CDCl_3_), were purchased from Sigma-Aldrich (St. Louis, MO). Hoechst 33258 solution, Dil Stain (1,1’-Dioctadecyl-3,3,3’,3’-Tetramethylindocarbocyanine Perchlorate (‘DiI’; DiIC18(3))) were purchased from ThermoFisher Scientific. Phosphate buffered saline (PBS), human plasma (HP), normal saline (0.9 NaCl), Dulbecco’s modified Eagle medium (DMEM), Roswell Park Memorial Institute (RPMI) 1640 Media, penicillin-streptomycin solution (10,000 units penicillin per ml and 10 mg streptomycin per ml), fetal bovine serum (FBS), 0.22 µm syringe filters, and Vybrant™ DiO Cell-Labeling Solution, Cell Counting Kit-8 (CCK-8) were purchased from Thermo Fisher Scientific. RAW 264.7 macrophages and IC21 cells were obtained from the American Type Culture Collection (ATCC). Preclinical Fenestra™ HDVC was purchased from MediLumine Inc. and OptiPrep™ Density Gradient Medium, 60 (w/v) iodixanol in water was purchased from Sigma-Aldrich (St. Louis, MO).

### Methods

#### Synthesis of 2,4,6-triiodophenyl oleate (TIPhO)

The synthetic procedures were performed as outlined in our previous works with a product yield of 89% [25]. ^1^HNMR analysis was also conducted using the same methods as before [25]. The resulting ^1^HNMR data can be found in Supplementary **Fig S1**.

<u>TIPhO:</u> Colorless oil; ^1^H NMR (500 MHz, CDCl_3_): δ/ppm: 8.05 (s, 2H), 5.33 – 5.30 (m, 2H), 2.62 (t, *J* = 7.5 Hz, 2H), 2.27 (t, *J* = 7.5 Hz, 1H), 2.04 – 1.94 (m, 4H), 1.82 – 1.76 (m, 2H), 1.59 (t, *J* = 7.3 Hz, 1H), 1.47 – 1.41 (m, 2H), 1.33 – 1.18 (m, 16H), 0.86 (t, *J* = 6.9 Hz, 3H). ^13^C NMR (126 MHz, CDCl_3_) δ 174.08, 151.86, 146.96, 129.93, 129.69, 91.79, 91.35, 77.37, 77.11, 76.86, 34.05, 31.93, 29.78, 29.69, 29.54, 29.35, 29.33, 29.17, 29.15, 29.13, 29.09, 27.22, 27.16, 24.94, 22.70, 14.12.

#### Formulation of ILNEs

The ILNEs were produced using a low-energy spontaneous emulsification technique, as outlined previously [26]. The oil phase components, namely TIPhO and Kolliphor ELP surfactant, were typically thoroughly mixed by heating at 50–60 ⶬ and vortexing for 5 minutes until complete dissolution and miscibility and the emergence of a clear phase. Subsequently, 0.1 NaCl saline (the aqueous phase) was swiftly introduced to the mixture, followed immediately by vortexing for 2–3 minutes, resulting in the formation of an opaque colloidal nanosuspension of ILNEs. The formulations were designed to maintain a *surfactant / (surfactant + oil)* weight ratio (SOR) at 20–50 wt% and *(surfactant + oil) / (surfactant + oil + water)* weight ratio (SOWR) at 40 wt%. The resultant samples were then passed through a 0.22 µm syringe filter for sterilization and to eliminate any potential aggregates or junctions. These samples were subsequently stored either at room temperature (RT) or at 4 ⶬ. For comparative analysis, some samples underwent lyophilization, followed by rehydration and storage under the same conditions of RT or 4 ⶬ for further characterization.

#### NP size and zeta-potential measurements

Dynamic light scattering (DLS) analysis was conducted following the method outlined in previous studies [27]. Similarly, nanoparticle tracking analysis (NTA) by Zetaview was performed using a Zetaview PMX 110 V3.0 instrument (Particle Metrix GmbH, Germany), with data analysis conducted using the Zetaview NTA software as previously described [27].

#### Cryo-transmission electron microscopy (Cryo-TEM)

Cryo-grids containing ILNE3 samples were swiftly immersed in a 60:40 mixture of liquid ethane and propane using a FEI Vitrobot Mark IV (ThermoFisher Scientific) at 22°C and 95% humidity. Lacey carbon TEM grids were made hydrophilic via indirect O_2_/Ag plasma treatment with a PIE Scientific TergeoEM plasma cleaner. Each cryo-grid received 3 µL of ILNE3 sample in normal saline buffer on the carbon side of the TEM grid for 30 seconds, followed by blotting for 3-6 seconds with Whatman #595 filter paper, and then plunged into the ethane-propane mixture. The cryo-grids were then imaged under low-dose conditions using a 200 keV TFS Talos Arctica equipped with a Gatan K3 DED.

#### Ultraviolet-visible (UV-Vis) spectroscopy

Using a Cary 4000 spectrophotometer (Varian), UV-Vis absorption spectra were recorded in the range of 200–700 nm. Absorbance measurements were taken at the peak wavelengths (λ_max_) of 230 nm for CrEL surfactant and 244 nm for TIPhO compounds.

#### Viscosity measurements

Particle-tracking microrheology (PTMR) measures the mechanical properties of mucus at the length scale of its constituent biopolymers [28, 29]. The procedures were performed according to the reported method [30, 31].

#### Particle stability in serum and plasma

To assess the stability of NPs in FBS and HP, freshly prepared ILNE3 and ILNE4 in normal saline solution (0.9% NaCl) were separately incubated in FBS and HP at two concentrations (10% and 20% (v/v)). The samples were then placed in shaking incubators set at 200 rpm and 37ⶬ for over 48 hours. Size distribution and polydispersity were monitored using DLS at various time intervals (0, 1, 2, 4, 6, 24, and 48 hours).

#### Cell viability assessment

We conducted cell viability studies to assess the *in vitro* cytotoxicity of ILNE3 on RAW 264.7 and IC21 macrophage cells. Various concentrations of ILNE3 were prepared through serial dilution in full medium and applied to the cells. Before treatment, cells were cultured and seeded in a 96-well plate at a density of 10^4^ cells per well, allowing them to attach for 24 hours. Cell viability was assessed using the CCK-8 assay according to the manufacturer’s instructions, 48 hours after incubation with ILNE3. The data are presented as the mean ± standard deviation (SD) of six replicate wells.

#### Cellular uptake of ILNE3 in RAW264.7 macrophages

A 100,000 RAW 264.7 macrophage cells were seeded per well in an appropriate cell culture plate. A solution of ILNE3 (40 mg NPs/mL) containing 4 µL of 25 mM Dil dye in NP lipid cores was prepared. The 1 mL of ILNE3 solution underwent purification using a nab10 column to eliminate any free dye and excess ingredients, yielding 1.5 mL of purified ILNEs as a stock solution. NTA analysis revealed a concentration of 7.2 × 10^13^ particles/mL, with a ζ-potential of -12.8 mV, and a particle size of 72.9 ± 5.1 nm. Subsequently, two different concentrations of ILNE3, 10 µL and 20 µL aliquots, were separately added to 1 mL of DMEM. This resulted in concentrations of approximately 266 µg NPs/mL, Dil dye of ∼130 nM, and 7.2 × 10^11^ particles/mL for the lower concentration, and approximately 533 µg NPs/mL, Dil dye of 260 nM, and 14.4 × 10^11^ particles/mL for the higher concentration. The 100,000 cells were then incubated with these two respective ILNE3 concentrations for 4 and 24-hours. Following the incubation period, the media was removed, and the cells were washed twice with fresh warm DMEM (1 mL). Subsequently, 0.5 mL of Hoechst dye solution (2 µg/mL) was incubated with the cells for 20 minutes. The dye solution was then removed, and the cells were subjected to two additional washes with DMEM. Finally, 0.5 mL of DMEM was added to each well, and the cells were imaged using a fluorescence microscope.

#### Phantom Scan: Assessment of x-ray attenuation for quantification of iodine concentration

The samples were scanned using a Rigaku Quantum GX small-animal CT system. Imaging parameters included an x-ray voltage of 90 kVp, an anode current of 88 μA, and an exposure time of 120 ms per rotational step (total rotation: 360 degrees, 180 steps). Reconstructed images were generated on a 512 × 512 pixel grid with a pixel size of 66.7 μm × 66.7 μm. ILNE contrast agents were scanned alongside various concentrations of the clinical CT contrast agent Visipaque™ (iodixanol), with deionized water serving as the control (0 HU).

#### Animal studies

All animal procedures adhered to the guidelines set by the Institutional Animal Care and Use Committee (IACUC) at the University of North Carolina at Chapel Hill. Healthy male C57BL/6 mice, aged 6-8 weeks, were sourced from Jackson Laboratory. The mice (five per cage) were randomly assigned to control and contrast agent groups. Animals were anesthetized and intubated before any procedure with the assistance of the Division of Comparative Medicine (DCM). All non-imaging procedures were performed by DCM staff.

Sequential imaging was performed on the Quantum GX2 micro-CT scanner which allows rapid, repeated scans of the same region. A standard abdominal imaging protocol was utilized with isoflurane anesthesia. We used this approach to replicate the four-phase liver CT imaging approach on clinical scanners. Images were acquired during the administration of the ILNE contrast agent at total iodine doses of ∼300 mgI/kg. Dynamic images were acquired every 5 seconds for 60 seconds during which a tail vein injection of the contrast agent will be administered. Repeat single-phase scans were subsequently at 5 min, 1, 2-, 4-, 24- and 72-hours post-injection. For this study, healthy mice were utilized. A reference sample of the contrast agent was scanned within the field-of-view of the liver to provide an external reference for the degree of iodine attenuation.

Analysis: Regions of interest (ROIs) were drawn on the kidney, spleen, liver, aorta, and heart to evaluate the attenuation of the injected contrast agent in Hounsfield Units (HU). We evaluated the relative hepatic uptake of the contrast agent as measured by increased attenuation.

The dose-escalation toxicity study involved the administration of a single dose of 0.9% saline solution and freshly prepared contrast agent formulations at concentrations of 300 mgI/mL (equivalent to 800 mg ILNEs/mL) and 750 mgI/mL (equivalent to 2000 mg ILNEs/mL) in 0.9% saline solution via tail vein intravenous (i.v.) injection. The general behavior and changes in body weight of the mice were monitored and recorded every other day for up to 10 days.

#### Hematology and blood chemistry studies

Blood samples were collected from C57BL/6 mice at three and ten days post-ILNE3 injection. Using 25 G syringes, blood was drawn via cardiac puncture into K2-EDTA microtainer tubes (500 µL; BD 365974, USA). Cell blood count and hematology parameters were measured with an IDEXX Procyte DX (Westbrook, Maine, USA). Plasma was separated by centrifuging the blood samples at 2,000 x G for 15 minutes, and blood chemistry parameters were analyzed using an Alfa Wassermann Vet Axcel (West Caldwell, NJ, USA) blood chemistry analyzer.

#### Histopathological assessment

Three days post-ILNE3 injection, tissues (liver, spleen, kidney, heart, and lung) were collected and preserved in 10% neutral buffered formalin (Sigma-Aldrich) for histological examination. The histological analysis was conducted by the Center for Gastrointestinal Biology and Disease Core at UNC, following previously described methods [32].

#### Porcine animal model

Imaging of a 16.6 kg porcine was performed on a clinical CT scanner (Siemens Force) using an abdominal protocol at 120kVp and mean x-ray tube current of 32mA. Images were reconstructed with a standard soft tissue kernel. All studies were performed under IACUC approval and with the assistance of DCM veterinarians. Imaging was acquired at 2 hours after the i.v. administration of the ILNE3 contrast agent (300 mgI/mL). Iohexol (300 mgI/mL) based imaging was performed in a different animal for reference prior to and after the i.v. administration of contrast agent at 1.5 mL/kg. Imaging was performed during the arterial phase of contrast administration. Vital signs were continuously monitored per protocol. No acute changes in the animal’s vital signs were observed during the administration of contrast or during the scan

## Results

### Preparation and characterization of ILNE CT contrast agents

We synthesized an iodinated lipid compound, TIPhO, achieving an 89% yield through a one-step esterification reaction using EDC•HCl and DMAP (**Fig 1a**). The chemical structure was confirmed via _1_HNMR and ^13^CNMR (supplementary **Fig S1**), consistent with our previous reports [25].

**Fig 1.**
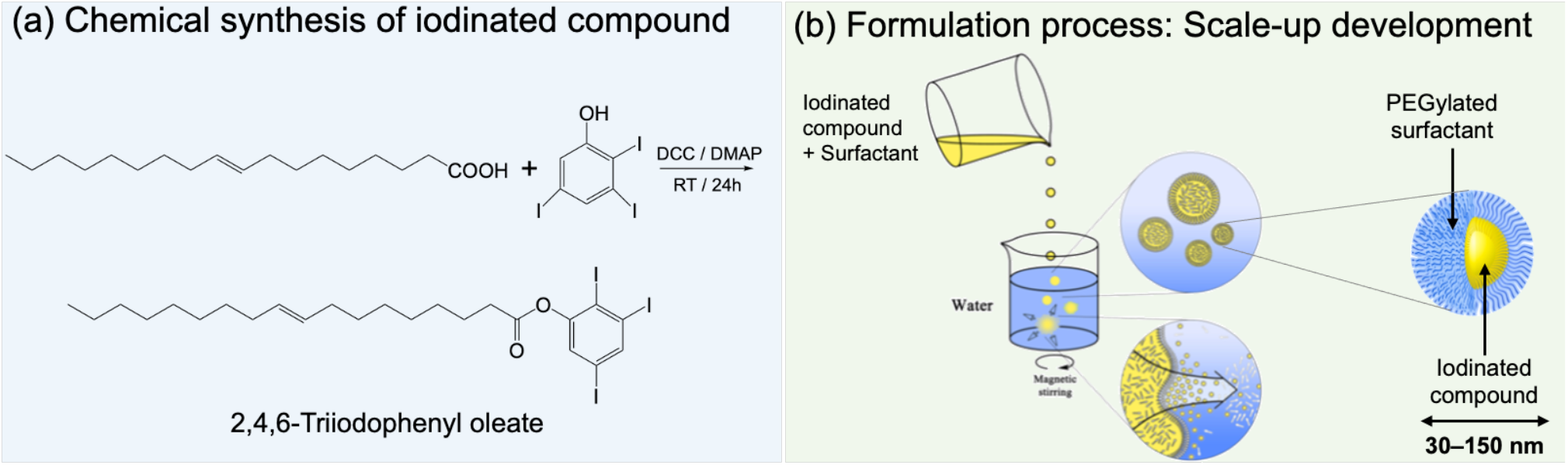
Preparation of ILNEs CT contrast agent. (a) Chemical synthesis of TIPhO via a one-step esterification reaction. (b) The formulation of ILNEs by a low-energy spontaneous emulsification process produced stable nanoemulsion droplets in the 30-150 nm size range, depending on the SOR ratio. Higher concentrations of the iodinated compound resulted in larger droplet sizes.

Using a green, scalable process, we employed a low-energy spontaneous emulsification technique without organic solvents or high-energy inputs. This involved mixing the iodinated compound (TIPhO) and the nonionic amphiphilic PEGylated surfactant Kolliphor^®^ ELP (CrEL) to form the oil phase at a SOR of 20– 50 wt%. The aqueous phase (0.9% normal saline) was then added at a SOWR of 40 wt% (**Fig 1b**). This produced stable, homogeneous formulations that remained stable for months to years at RT or 4ⶬ. The surfactant reduced interfacial tension and prevented nanodroplet aggregation, resulting in a stable nanoformulation.

#### ILNE Formulations

Four ILNE formulations (ILNE1–4) with iodinated cores and surface PEG molecules were designed with varying SOR ratios, affecting hydrodynamic size, ζ-potential, and iodine content (**Fig 2a, 2b**). SOR30 ILNEs were optimized for injectability, viscosity, and achieving a clinical dose of about 300 mgI/kg.

**Fig 2.**
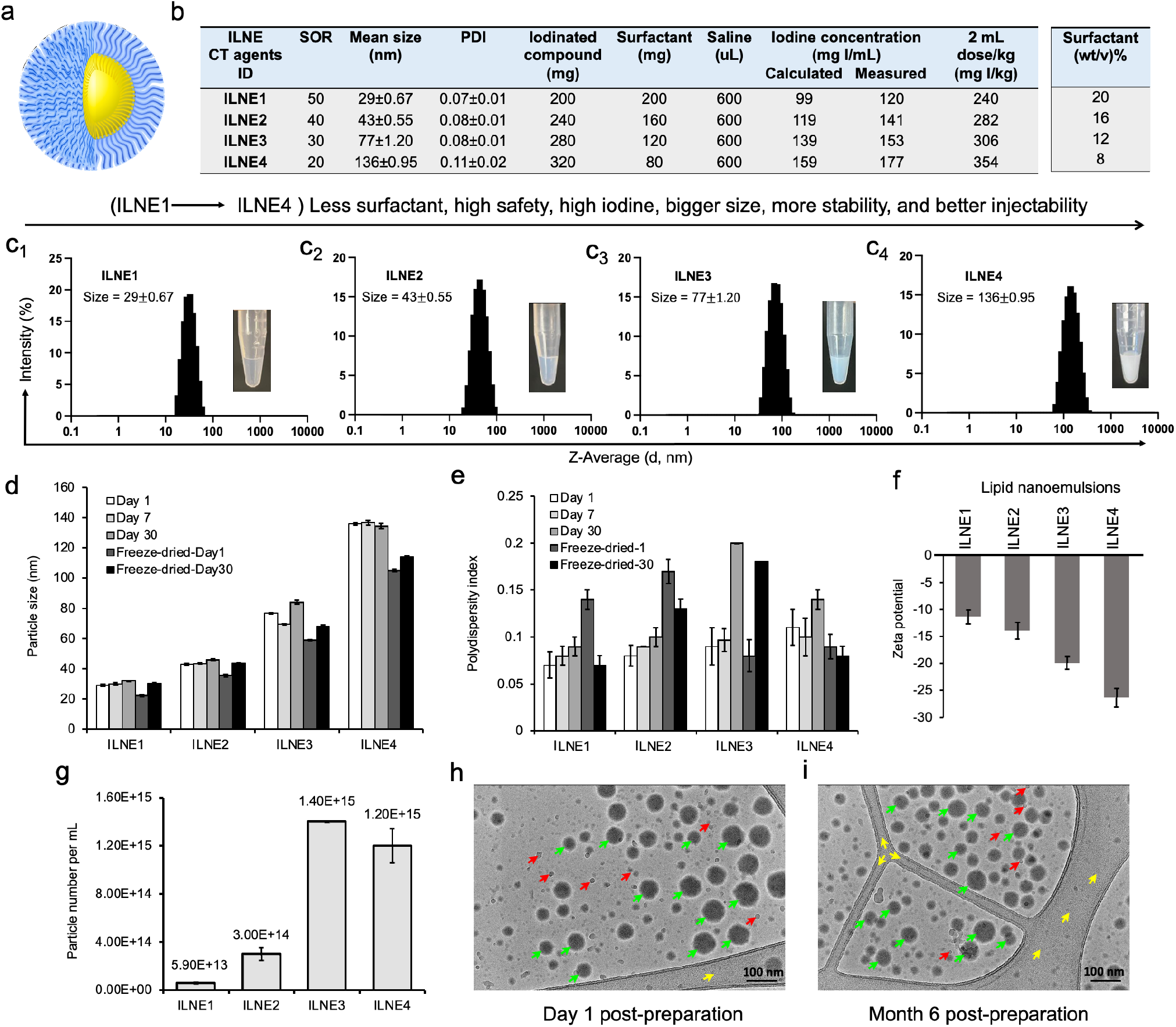
Characterization of the formulated lipid nanoemulsions. **(a)** Schematic representation of ILNEs composed of an oily core (yellow) surrounded by a PEG shell (blue). **(b)** Table showing the composition and ratio of all ingredients involved in the ILNE formulations (ILNE1–4) at SOWR 40%, and the final iodine concentrations of the injected dose. **(c1-c4)** Appearance and DLS histograms showing the size distribution of the four ILNEs. **(d, e)** Particle size and PDI over time measured by DLS. Samples were kept in aqueous dispersion at RT for different time intervals. Freeze-dried-1 and freeze-dried-30 are the redispersed lyophilized samples after one and 30 days of preparation, respectively. **(f)** Particle ζ-potential measured by Malvern Zetasizer. **(g)** Particle concentration of ILNEs determined by NTA (n=3). **(h, i)** Cryo-TEM images of ILNE3 at one day (h) and six months (i) post-preparation at RT. The green arrow indicates ILNE3 particles, the red arrow indicates ice particles, and the yellow arrow indicates the grid membrane.

#### Challenges and Solutions

Formulating efficient nanodroplet emulsions of lipidic compounds with free active functional groups (COOH or NH_2_) posed challenges [23, 25]. Blocking these groups improved homogeneity with surfactants enhancing formulation efficiency. Unmodified oleic acid formed highly viscous nanogels at lower concentrations (SOR 15–20 wt%) and solidified at higher concentrations (SOR > 20 wt%), resulting in less efficient nanoformulations (supplementary **Fig S2**). ILNE1–4 produced narrowly distributed sizes (30–135 nm) with low PDIs (<0.1), depending on the SOR ratio (**Fig 2c1–c4**). Sizes remained stable after 30 days, with slight size reduction in lyophilized samples (**Fig 2d**). Polydispersity was consistently <0.2, indicating high uniformity and stability (Fig 2e). Sizes and PDIs reduced after freeze-drying (supplementary **Fig S3**). Surface charge, determined by DLS, showed elevated negative ζ-potential with increasing TIPhO concentrations, ranging from –11.3 ± –1.32 to –26.3 ± –1.78 mV (**Fig 2f**).

#### Particle Concentration and Stability

Particle concentration for ILNE contrast agents, determined by Zetaview NTA analysis, revealed distinct particle numbers based on hydrodynamic size and SOR ratio: ILNE3 > ILNE4 > ILNE2 > ILNE1 with concentrations of 1.40E+15, 1.20E+15, 3.00E+14, and 5.9E+13 particles/mL, respectively (**Fig 2g**). ILNE3 was selected as the ideal contrast agent based on stability, viscosity, contrast efficiency, and clinical dose. Cryo-TEM images of ILNE3 showed similar morphology, size, and homogeneity for fresh and 6-month post-preparation samples (**Fig 2h**,**i**), consistent with DLS data.

#### Batch Reproducibility and Shelf-Life Stability

ILNE3 was assessed across different batch scales (2 mL, n=7 and 35 mL, n=1), revealing exceptional consistency (supplementary **Fig S4**). Particle sizes centered around 76.6 ± 1.2 nm with a low PDI of 0.08 ± 0.01. Shelf-life stability of ILNE3 and ILNE4 over a year at room temperature showed no significant size alterations, maintaining sizes of 81.1 ± 1.1 and 137 ± 3.8 nm, respectively, with PDIs < 0.2 (supplementary **Fig S5**). ILNE3 size and PDI also remained stable over six months at 4ⶬ (supplementary **Fig S6**).

#### Comparison with Commercial Agents

A commercial preclinical blood pool CT agent, Fenestra™ HDVC, exhibited a larger size (98 ± 0.62 nm) and higher PDI (0.37 ± 0.01) compared to ILNE3, which had better clarity, homogeneity, injectability, and a higher iodine payload (supplementary **Fig S7**).

#### Alternative Surfactants

Solutol^®^ HS15 (Kolliphor^®^ HS 15) was assessed as a replacement for CrEL in ILNE formulations, achieving similar iodine concentrations (∼353 mgI/kg) but larger particle sizes compared to CrEL-based formulations (supplementary **Table S1a**). Formulations with CrEL:Solutol^®^ HS15 (50:50) had sizes comparable to CrEL-based formulations (supplementary **Table S1b**). Our formulations contained up to 91.2-fold less CrEL excipients compared to chemotherapeutic paclitaxel (CrEL-paclitaxel) formulations (supplementary **Table S2**). Injected volume doses of ILNE3 across different body weights are presented in supplementary **Table S3**.

#### Nanoparticle Stability

Nanoparticles degrade due to chemical instability, pH sensitivity, aggregation, and interactions with biomacromolecules, affecting their efficacy. Ensuring NP stability in serum proteins is crucial. We evaluated ILNE3 and ILNE4 stability in human plasma (HP) and fetal bovine serum (FBS) over 48 hours. Two concentrations (10% and 20% v/v) of ILNEs in HP and FBS were incubated at 200 rpm and 37ⶬ. Particle size and polydispersity index (PDI) were monitored using dynamic light scattering (DLS). Results showed consistent sizes and PDIs in FBS, with a slight size increase in HP at 10% v/v after 24 hours, indicating good stability in both environments (**Fig 3a–d**).

**Fig 3.**
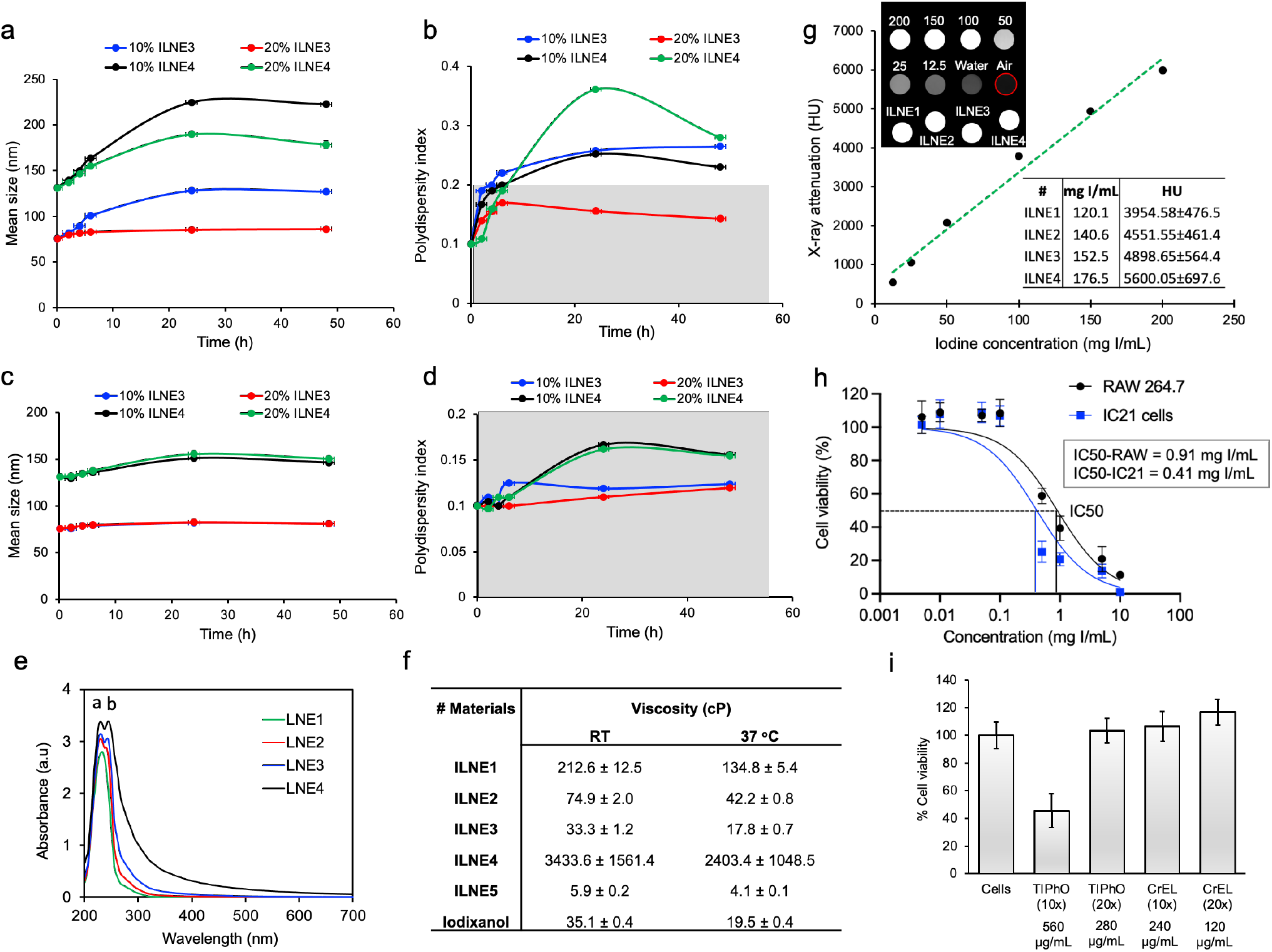
Characterization and stability of ILNEs in various conditions. (a-b) Particle size stability in HP and (c, d) FBS: 10% and 20% (v/v) ILNEs were incubated with (a, b) HP or FBS (c, d) for 48 hours. Particle size (a,c) and PDI (b,d) were measured at different time points using DLS. **(e)** UV-vis absorbance measurements of ILNE1–4, showing the peaks of CrEL and TIPhO at λmax of ∼230 nm and ∼244 nm, respectively. **(f)** Viscosity measurements of ILNE1–4 and ILNE5 (a 2x dilution of ILNE3), compared with iodixanol stock solution (300 mgI/mL) at 60% (wt/v). **(g)** Phantom scan of ILNEs: In vitro evaluation of x-ray attenuation properties (in HU) of ILNE1–4 using a calibration curve made by various dilutions of the iodixanol injection as a reference (expressed in mgI/mL). Air and water were used for normalization. Top view of Eppendorf tubes and source data are provided as a Source Data file. **(h)** CCK-8 cytotoxicity assay of immune cells: Macrophages RAW264.7 and IC21 cells were incubated with ILNE3 for 48 hours. Data are presented as the average ± SD (n = 6). **(i)** Cell viability studies of ILNE components: Two concentrations of CrEL and TIPhO were incubated with RAW 264.7 cells for 24 hours.

#### UV-vis Absorbance

Analysis of ILNE1–4 at a 3200x dilution revealed maximum wavelengths (λmax) peaks at 230 nm and 244 nm, confirming the presence of CrEL and TIPhO, respectively (**Fig 3e**). Absorbance spectra at various dilutions (1600x, 3200x, 6400x, and 12800x) were consistent (supplementary **Fig S8**).

#### Viscosity

Viscosity is important for designing injectable products. We compared the viscosity of all ILNE formulations with 2x diluted ILNE3 and clinical CT contrast agent iodixanol (Visipaque™) (300 mgI/mL) at 60% wt/v using PTMR. ILNE3 had lower viscosity than other ILNEs, measuring 33.3 ± 1.2 and 17.8 ± 0.7 cP at 25ⶬ and 37ⶬ, respectively, comparable to iodixanol. Diluting ILNE3 2x reduced its viscosity to 5.9 ± 0.2 and 4.1 ± 0.1 cP at 25ⶬ and 37ⶬ, respectively. ILNE3 also had lower viscosity than clinical iohexol (300 mgI/mL) and preclinical Fenestra™ HDVC products (**Fig 3f**).

#### Iodine Content

Iodine content is crucial for CT contrast agents. Most current hydrophilic CT contrast agents have iodine contents below 50%, such as iodixanol (49.1 wt%), iohexol (46.4 wt%), and iopromide (48.1 wt%). Our ILNEs, with 51.7 wt% iodine, show excellent x-ray attenuation. The CT value, measured in Hounsfield units (HU), increased linearly with iodine concentration in ILNE suspensions (supplementary **Fig S9**). We quantified iodine concentration using x-ray imaging and a calibration curve with iodixanol dilutions as a reference (**Fig 3g**). Data showed slightly higher x-ray attenuation for ILNEs compared to theoretical values (**Fig 2b**). Iodine concentrations ranged from 120 to 176 mgI/mL, providing 3955 to 5600 HU, depending on SOR ratios. Lower SOR values resulted in higher iodine content and contrast. ILNE3 at SOR30 and ILNE4 at SOR20 are promising for clinical CT imaging, allowing low-volume dosages (∼1.5–2 mL/kg) with iodine amounts aligning with clinical doses (∼300 mgI/mL). ILNE tubes showed similar contrast enhancement to 200 mgI/mL iodixanol, ensuring high contrast (**Fig 3g**).

#### In Vitro Toxicity

We assessed the in vitro toxicity of ILNE3 suspensions using the CCK-8 test after a 48-hour incubation with RAW 264.7 macrophages and IC21 immune cells at various concentrations (**Fig 3h**). The data revealed lower cytotoxicity for RAW macrophages compared to IC21 immune cells, with LC50 values of 0.91 and 0.41 mgI/mL, respectively. Given that ILNE3 suspensions are administered at 40 µL per mouse (20g), containing nearly 2 mL of blood, the injected dose amounts to 8 mg NPs/mL of mouse blood (equivalent to ∼2.9 mgI/mL mouse blood), as iodine constitutes approximately 36% of the final NPs. In cytotoxicity studies, the calculated concentration of NPs for in vitro cell incubation is considered to be within 100-fold less than that administered in animals. As shown in **Fig 3h**, concentrations up to 0.1 mgI/mL demonstrated 100% cell viability in both immune cell types, representing a 29-fold lower concentration than the injected dose in mice, indicating excellent biocompatibility. Moreover, the IC50 values for RAW 264.7 and IC21 cells were 11- and 5-fold less than the administered dose in mice, respectively. These results are notable, considering the CCK-8 assay conditions are more stringent than in vivo. During the 48-hour incubation in vitro, cells are in prolonged contact with ILNEs and free surfactants, whereas these surfactants are generally rapidly cleared from the bloodstream in vivo.

We also conducted separate cell viability studies on the CrEL and TIPhO compounds, which constitute ILNE3 (**Fig 3i**). Following a 24-hour incubation with RAW macrophages, full cell viability was maintained at concentrations of 240 and 280 µg/mL for CrEL and TIPhO, respectively—these concentrations represent a 20-fold reduction of the injected dose in mice. Upon increasing both concentrations 10-fold, CrEL did not adversely affect the cells, while TIPhO led to approximately 50% cell death among macrophages. Collectively, ILNE3 and its constituents demonstrated exceptional cytocompatibility in vitro relative to the injected dose in animals.

### In Vitro Cellular Uptake Study

Understanding nanoparticle uptake by cells is crucial for evaluating toxicity and biocompatibility. Inadequate internalization or cytotoxic effects can compromise cellular function. NPs must overcome biological barriers like the cell membrane and endosomal/lysosomal compartments to deliver therapeutic or contrast agents effectively. We prepared two concentrations of ILNE3 (266 and 533 µg/mL), encapsulating Dil lipophilic dye within their lipidic core, to monitor their internalization into RAW 264.7 macrophages (**Fig 4**). These samples were incubated with the cells for 4 and 24 hours. After incubation, the cells were washed, and the nuclei were stained with Hoechst dye before imaging. The images (**Fig 4**) showed that the higher concentration of particles exhibited slightly greater uptake after 4 hours. However, at both concentrations, ILNE3 displayed similar uptake intensity at 24 hours, significantly higher than at 4 hours. Overall, ILNE3, with a size of about 72.9 ± 5.1 nm and a ζ-potential of -12.8 mV (measured by NTA-zetaview), demonstrated remarkable internalization within the cytoplasm of macrophage cells, indicating their potential for targeting and biocompatibility.

**Fig 4.**
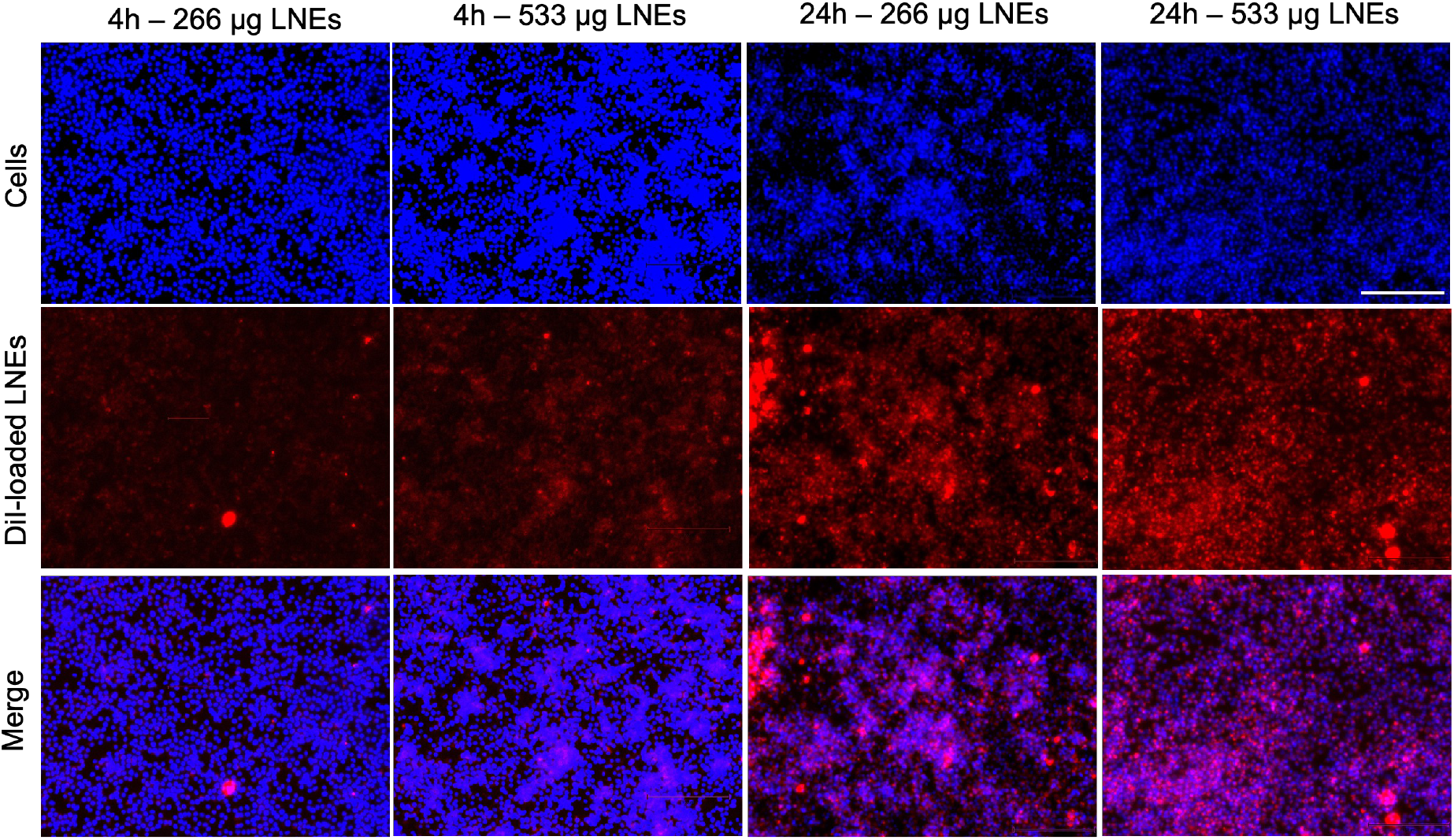
Cellular uptake study of Dil dye-loaded ILNE3 in RAW 264.7 macrophage cells. Two concentrations of ILNE3 were incubated in the cells, and the internalization of particles was observed using an optical fluorescence microscope at 4 and 24-hour time points. Images in each column (top to down) show: Hoechst dye (blue) for staining cell nucleus, Dil dye-labeled ILNE3 (red), and overlapping (blue-red). The scale bar indicates 150 µm.

### In Vivo CT Contrast and Pharmacokinetics in Mice

We administered ILNE3 intravenously at 2 mL/kg to C57BL/6 mice. Enhanced contrast was observed in the heart and liver compared to pre-injection images, evident in both coronal and transverse views (**Fig 5, top**). This confirms the rapid distribution of ILNE3 throughout the vasculature, highlighting the major veins of the heart ventricles, aorta, and liver. Over 72 hours, no clinical signs of disorder or toxicity were observed. ILNE3 consistently exhibited enhanced contrast, distribution, and clearance behavior. The images revealed a gradual increase in contrast in the heart and liver up to 4 hours post-injection. By 24 hours, ILNE3 had been entirely cleared from the bloodstream, with no detectable signals, while notably accumulating in the liver, resulting in pronounced contrast. By 72 hours, CT signals in the liver were significantly reduced, confirming elimination from the body. ILNE3 functions as a blood pool CT contrast agent (BPCAs), boasting a half-life of at least 4 hours, prolonging the scan window, and subsequently undergoing hepatic metabolism, circumventing renal clearance. It is effectively eliminated from the body within approximately 3 days post-injection. These attributes match the criteria for optimal BPCAs in clinical settings, featuring a kidney-safe formulation with improved contrast enhancement, followed by rapid clearance from the body post-scanning.

**Fig 5.**
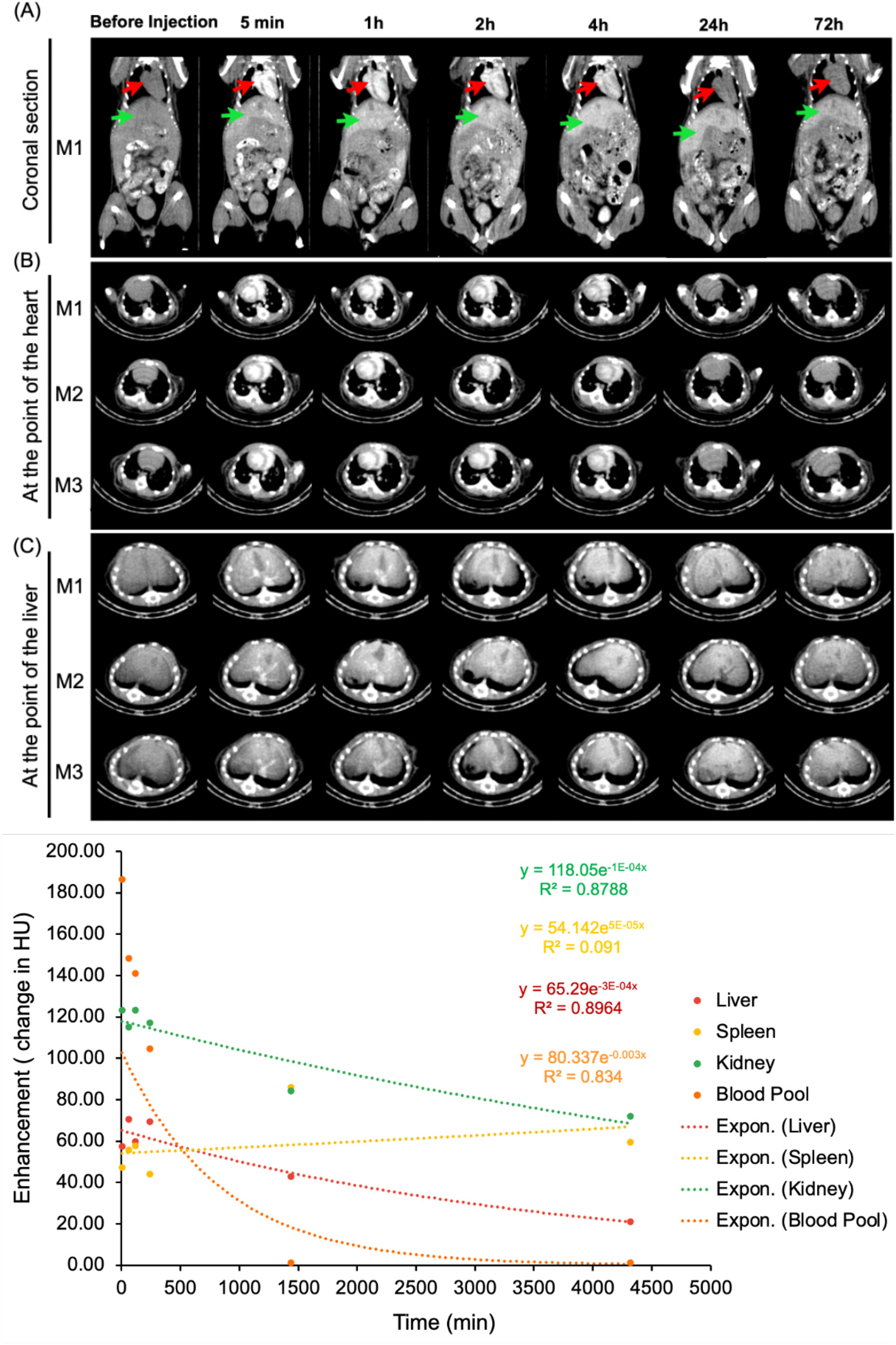
*In vivo* micro-CT imaging, longitudinal studies of biodistribution after i.v. administration of ILNE3 in C57BL/6 mice. Mice IDs are M1–M3 (n=3, biologically independent samples). (Top) Maximum intensity projection at representative times: (A) coronal view of mice pre- and post-injection at different time points. The red arrow represents the heart, and the green arrow represents the liver. (B) Transverse section at the view of the heart. (C) Transverse section at the view of the liver. (bottom) Quantitative analysis of the x-ray attenuation values in HUs where the ROIs were placed in the heart, liver, spleen, and kidneys.

Contrast enhancement was measured in the blood pool (left ventricle ROI), liver, spleen, and kidneys over three days (**Fig 5, bottom**). Blood pool enhancement peaks at the time of injection, achieving nearly 200 HU relative to the background. The kidney parenchyma shows expected vascular enhancement, with no significant contrast accumulation in the collecting system despite parenchymal enhancement. Liver enhancement peaks hours after injection as the agent accumulates in the liver. There is some renal excretion of iodine due to the ultrasmall particles within the ILNE3, with the bladder showing peak enhancement around the 4-hour mark based on mouse images (**Fig 5, top**).

### In Vivo Biocompatibility of ILNE3

To ensure the safety of ILNE3 as a CT contrast agent, we conducted two experiments on C57BL/6 mice.

#### Experiment (i)

A control group (n=2) received saline, and a treatment group (n=3) received ILNE3 at 300 mgI/kg. Three days post-injection and CT scanning, both groups were sacrificed for blood analysis and histological examination of major organs (heart, liver, kidney, lung, and spleen). Analysis (**Fig 6a**) showed no significant differences in liver/kidney functions or complete blood count (CBC) between the groups. Macroscopic examination and H&E stains indicated no toxicity, necrosis, or pathological alterations (**Fig 6b**), confirming ILNE3’s safety.

**Fig 6.**
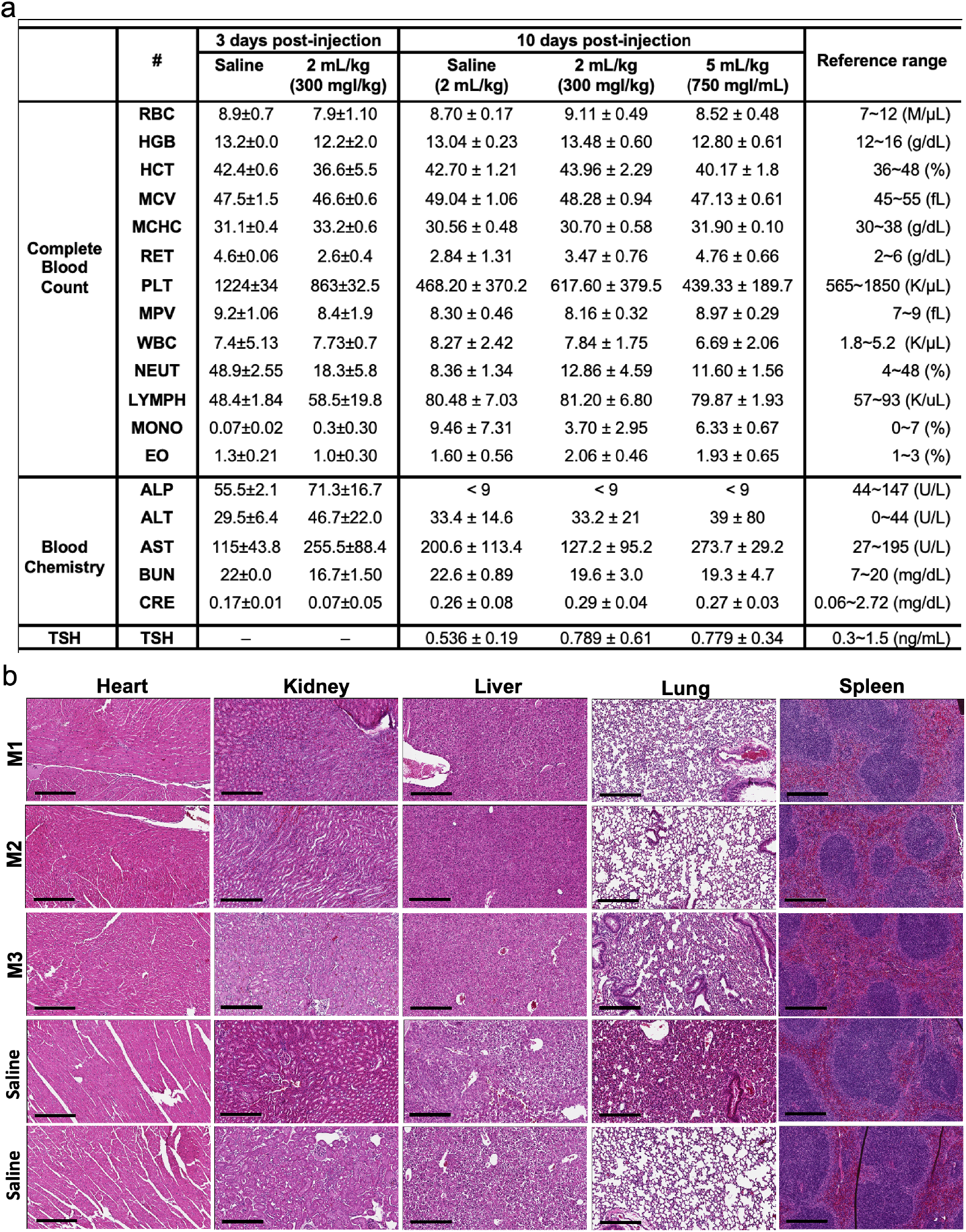
Blood analysis and histological examination post-ILNE3 injection. **(a)** Blood analysis and liver/kidney function tests were conducted at two distinct time points: three days following the injection of saline (n=3 biologically independent mice) and ILNE3 (2 mL/kg, n=3 biologically independent mice), and ten days post-injection of saline (2 mL/kg, n=5), ILNE3 (2 mL/kg, n=5), and ILNE3 (5 mL/kg, n=3). Thyroid-stimulating hormone (TSH) analysis was also determined at 10 days post-injection. Data are presented as Mean ± SD. Abbreviations: RBC (red blood cells), HGB (hemoglobin), HCT (hematocrit), MCV (mean corpuscular volume), MCHC (mean corpuscular hemoglobin concentration), RET (reticulocytes), PLT (platelets), MPV (mean platelet volume), WBC (white blood cells), NEUT (neutrophils), LYMPH (lymphocytes), MONO (monocytes), EO (eosinophils), ALP (alkaline phosphatase), ALT (alanine aminotransferase), AST (aspartate aminotransferase), BUN (blood urea nitrogen), CERA (creatinine). **(b)** H&E staining histology of organs (heart, kidney, liver, lung, and spleen) harvested three days post-injection of ILNE3 at 2 mL/kg (equivalent to 800 mg ILNE3/kg, n=3) compared with saline-injected mice (n=2). (M1–M3) are mice IDs.

### Experiment (ii)

Three groups were compared: saline-injected mice (n=5), ILNE3-injected mice at 300 mgI/kg (n=5), and ILNE3-injected mice at 750 mgI/kg (n=3). After 10 days, CBC, blood chemistry, and TSH levels were analyzed. Data showed no significant weight changes (supplementary **Fig S10**) or differences in blood analysis and liver/kidney function tests (**Fig 6a**). AST levels slightly elevated three days post-injection but normalized after 10 days. The higher dose of 750 mgI/kg showed only a minor AST increase, suggesting ILNEs accumulate in the liver before dissipating. These findings support ILNE3’s exemplary biosafety for translational studies.

### In Vivo CT Contrast in Porcine Model

To further verify scalability and safety, we injected ILNE3 intravenously into a 16.6 kg porcine at 300 mg/kg (∼2 mL/kg) and compared it to clinical iohexol at a similar dose, using the pre-contrast state as a baseline. For iohexol, we scanned the porcine at 30 seconds (arterial phase) due to rapid renal clearance. Improved contrast was observed in the heart (**Fig 7**, middle column), followed by significant kidney contrast. In contrast, ILNE3-injected porcine scanned 1-hour post-injection (**Fig 7**, right column) showed heart visualization and enhanced liver contrast with minimal kidney imaging. This suggests ILNE3’s potential as a kidney-safe BPCA. A 3D reconstructed image (**Fig 8**) further illustrates clear delineation of heart chambers, arterial and venous vasculature, and hepatic vessels.

**Fig 7.**
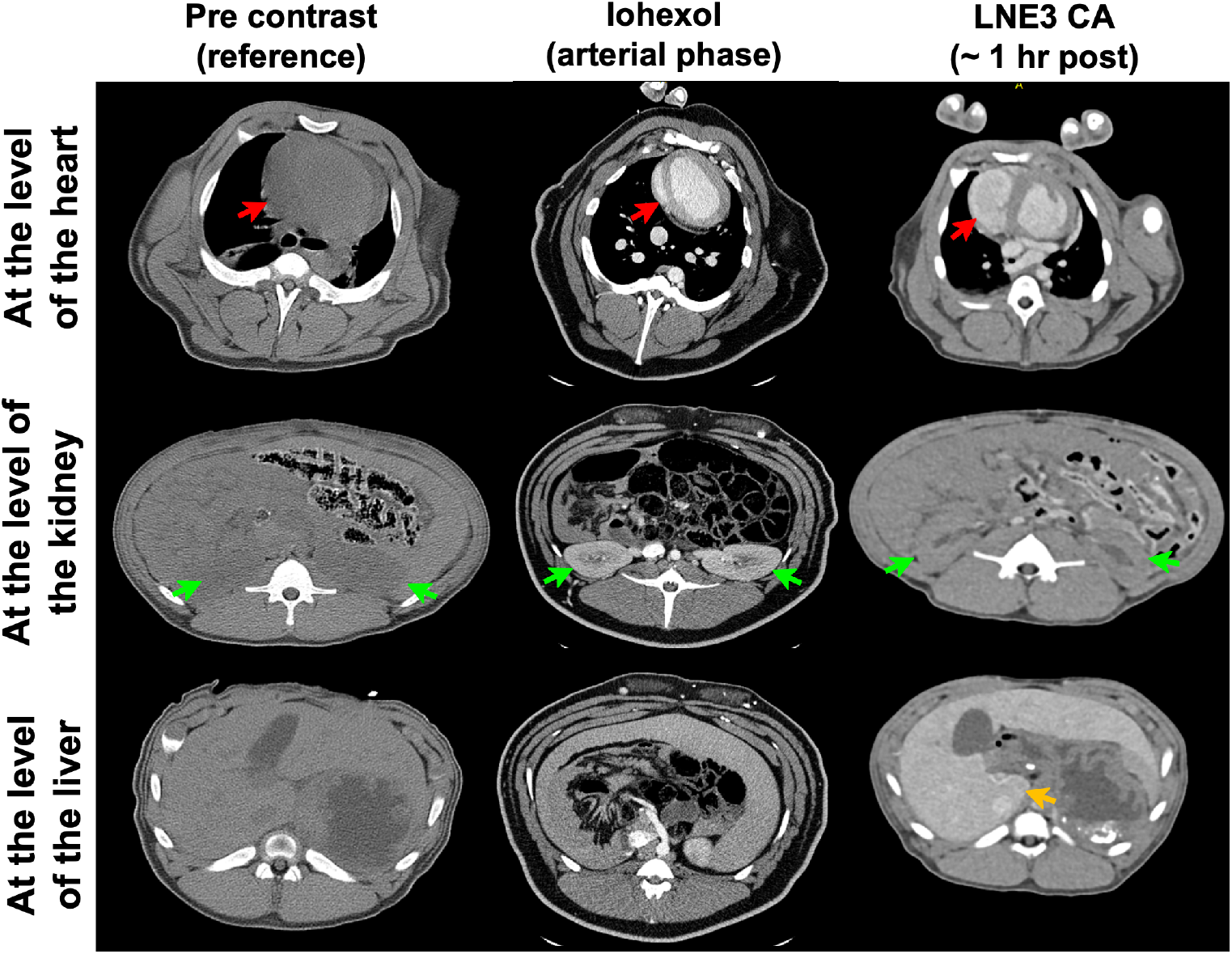
In vivo CT imaging of a porcine animal model injected with ILNE3 at a dose of 300 mgI/mL compared to a similar dose of iohexol. Imaging of the iohexol was acquired during the arterial phase, at 30 seconds post-injection. ILNE3 imaging was performed at approximately 1 hour post injection due to our workflow. Notably, the heart, kidney, and liver were identified and indicated by red, green, and yellow arrows, respectively, facilitating clear visualization and comparison of contrast enhancement dynamics between the two contrast agents. Significant arterial enhancement remains at the 1-hour time point with the ILNE3 imaging, despite the significant delay in imaging. Increased liver enhancement is also seen, due to the liver uptake.

**Fig 8.**
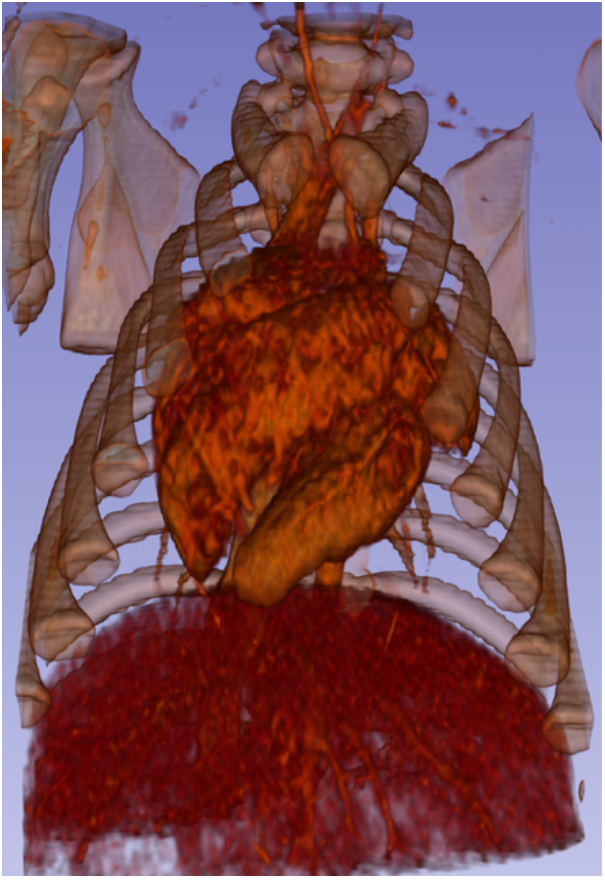
3D Rendering of porcine post-ILNE3 injection. 3D rendering image of a porcine subject 1-hour post-injection of ILNE3 contrast agent at a dose of 300 mgI/kg. The image shows clear delineation of the heart chambers, arterial and venous vasculature, and hepatic vessels. Diffuse liver uptake is also observed at this time point.

## Discussion

Nano-CT contrast agents are primarily used in preclinical studies involving polymers [33, 34], liposomes [35, 36], and metallic materials [37]. Our approach focuses on engineering safe, scalable intravenous iodinated lipid nanodroplet emulsion (ILNE) CT contrast agents. These agents offer significant contrast and minimal toxicity for clinical use, addressing the shortcomings of current clinical and NP-based agents due to their excellent biocompatibility, high iodine payloads, and robust CT intensity signals. Additionally, their surface can be easily labeled with targeted antibodies, enhancing CT imaging specificity and sensitivity. Our ∼75 nm-sized ILNEs enable extended blood circulation, expanding the scan window and allowing slower injection rates, minimizing infusion reactions and avoiding renal excretion, thus reducing kidney injury risks.

Previous iodinated contrast agents developed by our team and others suffer from long-term stability issues and excessive surfactant content (50-70 wt%) [26, 38]. This impedes clinical development due to toxicity from excessive excipients and the need for higher doses for effective contrast. These formulations also show prolonged accumulation in animals, increasing toxicity risks [26, 38].

The chemical structure and composition ratio between surfactant and iodinated compounds in nanoformulations significantly affect size, surface charge, iodine content, blood circulation time, pharmacokinetics, and biodistribution [25, 26, 38]. Hydrophilic small molecules are rapidly taken up by the kidneys, potentially causing kidney dysfunction, akin to clinical CT agents. Conversely, nanosized formulations (20–200 nm) undergo hepatic pathways and preferentially accumulate in tumor vasculature due to the enhanced permeability and retention (EPR) effect, aligning with our contrast agents.

Our spontaneous emulsification approach for producing lipid nanoemulsions saves time and money, reduces material degradation risk, and facilitates scalable production. This method ensures efficient size reduction with uniform distribution, enhancing bioavailability and shelf-life. It also enables sterile filtration with minimal clogging and transparent emulsion production, crucial for achieving stable and effective contrast agents.

Our prior efforts led to the synthesis of iodine-modified lipid compounds like cholecalciferol, α-tocopherol, and mono and triglyceride compounds as iodinated cores of successful preclinical CT imaging agents [25, 26, 38]. Notably, the commercialized Fenestra™ HDVC product by MediLumine is based on our iodinated α-tocopherol LNEs, achieving a two-fold increase in iodine concentration (100 mgI/mL) compared to its previous preclinical products. This allows researchers to use lower doses for vascular and hepatic imaging in small animals.

Our concept aims to reduce excipients while maximizing iodinated molecules to enhance safety and efficiency. Our findings indicate that a surfactant-to-oil ratio (SOR) of 20-30 wt% results in enhanced stability, reduced toxicity, and improved efficiency, aligning with the clinical dose of approximately 2 mL/kg (300 mgI/kg). These formulations accumulate in the liver for about three days before clearance, making them ideal CT contrast agents. The SOR ratio is key in varying physicochemical properties, toxicity, contrast enhancement, and clinical suitability.

Cytotoxicity assessments on RAW 264.7 macrophages and IC21 immune cells, and studies on C57BL/6 mice and porcine models, validate the safety and compatibility of our ILNEs CT agent. Histological analysis shows no necrosis or tissue damage in main organs. Comprehensive evaluations, including blood chemistry, CBC, and TSH analyses at 3 and 10 days post-injection, along with ongoing monitoring of animal behavior and weight, support the safety and potential efficacy of our ILNEs. Future investigations will focus on thorough toxicity assessments of the liver and thyroid glands across various species, facilitating the transition from preclinical research to clinical application. Our ILNEs can selectively target hepatocytes for functional and anatomical imaging, staging and monitoring fatty liver disease, and quantifying liver tumor burdens, validating their potential for heart and liver imaging.

## Conclusion

We introduce TIPhO lipid nanoemulsions (ILNEs) as a promising nano-contrast agent for clinical CT imaging, overcoming the limitations of traditional hydrophilic molecules and other nano-based materials. With a size of approximately 78 nm and minimal excipients, ILNEs are safe, scalable, and stable for intravenous use over several years. They follow the hepatic pathway and are eliminated within 72 hours post-injection. In vitro and in vivo studies in small and large animal models have validated the safety and contrast enhancement of ILNEs, showing superior performance compared to preclinical (Fenestra™ HDVC) and clinical (Omnipaque™) CT agents. Our preliminary results demonstrate excellent blood pool contrast with a half-life greater than 4 hours and enhanced liver imaging, with no observed pathology in the kidney, liver, or thyroid, and no adverse effects on blood function. These findings support the clinical translation of ILNEs, aiming to define the ideal nanoparticle format for safe and effective CT liver tumor imaging. We aim to assess this agent for detecting, characterizing, and monitoring treatment responses in primary and metastatic liver tumors. Ongoing non-GLP toxicology and pharmacokinetic studies in various animal species further ensure the safety of ILNEs, paving the way for their commercial use as a transformative CT contrast agent.

## Acknowledgments

MFA is supported by the Carolina Cancer Nanotechnology Training Program, funded by NCI grant T32CA196589. Research in the Kabanov lab received partial funding from NCI grant CA264488. The Whitehead lab is partially supported by NIH NIGMS grant 1P20GM146584-01. Special thanks to Lucas Plott at the Center for Esophageal Diseases & Swallowing (CEDS) at UNC-CH for assistance with viscosity measurements and Jon Frank at the Biomedical Research Imaging Center (BRIC) at UNC-CH for help with animal injections and other tasks. We used UNC-licensed Microsoft Bing Copilot to enhance the clarity and style of our phrasing while preserving originality, scientific content, and integrity of our research findings.

## Author contributions

**Conceptualization:** Mohamed F. Attia, Yueh Z. Lee

**Methodology:** Mohamed F. Attia, Ryan N. Marasco, Samuel Kwain, Yueh Z. Lee

**Investigation:** Mohamed F. Attia, Ryan N. Marasco, Samuel Kwain, Yueh Z. Lee

**Writing -Original Draft:** Mohamed F. Attia

**Writing -Review & Editing:** Mohamed F. Attia, Daniel C. Whitehead, Alexander Kabanov, Yueh Z. Lee

**Funding Acquisition:** Daniel C. Whitehead, Alexander Kabanov, Yueh Z. Lee

## References

[1] D. Xi, S. Dong, X. Meng, Q. Lu, L. Meng, and J. Ye, “Gold nanoparticles as computerized tomography (CT) contrast agents,” Rsc Advances, vol. 2, no. 33, pp. 12515–12524, 2012.

[2] W. E. Ghann, O. Aras, T. Fleiter, and M.-C. Daniel, “Syntheses and characterization of lisinopril-coated gold nanoparticles as highly stable targeted CT contrast agents in cardiovascular diseases,” Langmuir, vol. 28, no. 28, pp. 10398–10408, 2012.

[3] D. S. Gierada and K. T. Bae, “Gadolinium as a CT contrast agent: assessment in a porcine model,” Radiology, vol. 210, no. 3, pp. 829–834, 1999.

[4] L. Robison et al., “A bismuth metal–organic framework as a contrast agent for X-ray computed tomography,” ACS Applied Bio Materials, vol. 2, no. 3, pp. 1197–1203, 2019.

[5] N. Lee, S. H. Choi, and T. Hyeon, “Nano-sized CT contrast agents,” Advanced Materials, vol. 25, no. 19, pp. 2641–2660, 2013.

[6] M. Tepel, P. Aspelin, and N. Lameire, “Contrast-induced nephropathy: a clinical and evidence-based approach,” Circulation, vol. 113, no. 14, pp. 1799–1806, 2006.

[7] H. Y. Zhao et al., “Synthesis and application of strawberry-like Fe3O4-Au nanoparticles as CT-MR dual-modality contrast agents in accurate detection of the progressive liver disease,” Biomaterials, vol. 51, pp. 194–207, 2015.

[8] R. Cheheltani et al., “Tunable, biodegradable gold nanoparticles as contrast agents for computed tomography and photoacoustic imaging,” Biomaterials, vol. 102, pp. 87–97, 2016.

[9] A. C. Lohana, S. Neel, V. Deepak, and M. Schauer, “Intrathecal iodinated contrast-induced transient spinal shock,” BMJ Case Reports CP, vol. 13, no. 12, p. e237610, 2020.

[10] A. Kistner et al., “Negative effects of iodine-based contrast agent on renal function in patients with moderate reduced renal function hospitalized for COVID-19,” BMC nephrology, vol. 22, pp. 1–10, 2021.

[11] M. S. Davenport, S. Khalatbari, R. H. Cohan, J. R. Dillman, J. D. Myles, and J. H. Ellis, “Contrast material–induced nephrotoxicity and intravenous low-osmolality iodinated contrast material: risk stratification by using estimated glomerular filtration rate,” Radiology, vol. 268, no. 3, pp. 719–728, 2013.

[12] A.-L. Faucon, G. Bobrie, and O. Clément, “Nephrotoxicity of iodinated contrast media: From pathophysiology to prevention strategies,” European journal of radiology, vol. 116, pp. 231–241, 2019.

[13] J.-j. Fu et al., “Bismuth chelate as a contrast agent for X-ray computed tomography,” Journal of nanobiotechnology, vol. 18, pp. 1–10, 2020.

[14] L. Bolognese et al., “Impact of iso-osmolar versus low-osmolar contrast agents on contrast-induced nephropathy and tissue reperfusion in unselected patients with ST-segment elevation myocardial infarction undergoing primary percutaneous coronary intervention (from the Contrast Media and Nephrotoxicity Following Primary Angioplasty for Acute Myocardial Infarction [CONTRAST-AMI] Trial),” The American journal of cardiology, vol. 109, no. 1, pp. 67–74, 2012.

[15] T. C. Owens, N. Anton, and M. F. Attia, “CT and X-ray contrast agents: current clinical challenges and the future of contrast,” Acta Biomaterialia, 2023.

[16] X. Li, N. Anton, G. Zuber, and T. Vandamme, “Contrast agents for preclinical targeted X-ray imaging,” Advanced drug delivery reviews, vol. 76, pp. 116–133, 2014.

[17] A. S. Klymchenko, F. Liu, M. Collot, and N. Anton, “Dye-loaded nanoemulsions: biomimetic fluorescent nanocarriers for bioimaging and nanomedicine,” Advanced healthcare materials, vol. 10, no. 1, p. 2001289, 2021.

[18] J. A. Olzmann and P. Carvalho, “Dynamics and functions of lipid droplets,” Nature reviews Molecular cell biology, vol. 20, no. 3, pp. 137–155, 2019.

[19] A. Pol, S. P. Gross, and R. G. Parton, “Biogenesis of the multifunctional lipid droplet: Lipids, proteins, and sites,” Journal of Cell Biology, vol. 204, no. 5, pp. 635–646, 2014.

[20] R. Bouchaala et al., “Integrity of lipid nanocarriers in bloodstream and tumor quantified by near-infrared ratiometric FRET imaging in living mice,” Journal of controlled release, vol. 236, pp. 57–67, 2016.

[21] J. Choi et al., “Targeting tumors with cyclic RGD-conjugated lipid nanoparticles loaded with an IR780 NIR dye: In vitro and in vivo evaluation,” International Journal of Pharmaceutics, vol. 532, no. 2, pp. 677–685, 2017.

[22] A. Saito et al., “An azide-tethered Cremophor® ELP surfactant allowing facile post-surface functionalization of nanoemulsions,” Bulletin of the Chemical Society of Japan, vol. 93, no. 4, pp. 568–575, 2020.

[23] M. F. Attia, M. I. Swasy, R. Akasov, F. Alexis, and D. C. Whitehead, “Strategies for High Grafting Efficiency of Functional Ligands to Lipid Nanoemulsions for RGD-Mediated Targeting of Tumor Cells In Vitro,” ACS Applied Bio Materials, vol. 3, no. 8, pp. 5067–5079, 2020.

[24] M. F. Attia et al., “Functionalization of nano-emulsions with an amino-silica shell at the oil– water interface,” RSC advances, vol. 5, no. 91, pp. 74353–74361, 2015.

[25] M. F. Attia et al., “Functionalizing nanoemulsions with carboxylates: impact on the biodistribution and pharmacokinetics in mice,” Macromolecular Bioscience, vol. 17, no. 7, p. 1600471, 2017.

[26] M. F. Attia et al., “Biodistribution of X-ray iodinated contrast agent in nano-emulsions is controlled by the chemical nature of the oily core,” ACS nano, vol. 8, no. 10, pp. 10537–10550, 2014.

[27] M. F. Attia et al., “Enhancing drug delivery with supramolecular amphiphilic macrocycle nanoparticles: selective targeting of CDK4/6 inhibitor palbociclib to melanoma,” Biomaterials Science, vol. 12, no. 3, pp. 725–737, 2024.

[28] D. B. Hill et al., “A biophysical basis for mucus solids concentration as a candidate biomarker for airways disease,” PloS one, vol. 9, no. 2, p. e87681, 2014.

[29] S. K. Lai, Y.-Y. Wang, D. Wirtz, and J. Hanes, “Micro-and macrorheology of mucus,” Advanced drug delivery reviews, vol. 61, no. 2, pp. 86–100, 2009.

[30] T. G. Mason, “Estimating the viscoelastic moduli of complex fluids using the generalized Stokes–Einstein equation,” Rheologica acta, vol. 39, pp. 371–378, 2000.

[31] B. S. Schuster, J. S. Suk, G. F. Woodworth, and J. Hanes, “Nanoparticle diffusion in respiratory mucus from humans without lung disease,” Biomaterials, vol. 34, no. 13, pp. 3439–3446, 2013.

[32] C. Lim et al., “Enhancing CDK4/6 inhibitor therapy for medulloblastoma using nanoparticle delivery and scRNA-seq–guided combination with sapanisertib,” Science advances, vol. 8, no. 4, p. eabl5838, 2022.

[33] M. Yin et al., “Precisely translating computed tomography diagnosis accuracy into therapeutic intervention by a carbon-iodine conjugated polymer,” Nature Communications, vol. 13, no. 1, p. 2625, 2022.

[34] F. Hyafil et al., “Noninvasive detection of macrophages using a nanoparticulate contrast agent for computed tomography,” Nature medicine, vol. 13, no. 5, pp. 636–641, 2007.

[35] H. Xu et al., “Nanoliposomes co-encapsulating CT imaging contrast agent and photosensitizer for enhanced, imaging guided photodynamic therapy of cancer,” Theranostics, vol. 9, no. 5, p. 1323, 2019.

[36] K. B. Ghaghada, A. F. Sato, Z. A. Starosolski, J. Berg, and D. M. Vail, “Computed tomography imaging of solid tumors using a liposomal-iodine contrast agent in companion dogs with naturally occurring cancer,” PloS one, vol. 11, no. 3, p. e0152718, 2016.

[37] J. W. Lambert et al., “An intravascular tantalum oxide–based CT contrast agent: preclinical evaluation emulating overweight and obese patient size,” Radiology, vol. 289, no. 1, pp. 103–110, 2018.

[38] M. F. Attia, N. Anton, R. Akasov, M. Chiper, E. Markvicheva, and T. F. Vandamme, “Biodistribution and toxicity of X-ray iodinated contrast agent in nano-emulsions in function of their size,” Pharmaceutical research, vol. 33, pp. 603–614, 2016.

