## Supplementary material for "Toward the Clinical Translation of Safe Intravenous Long Circulating Iodinated Lipid Nanoemulsion Contrast Agents for CT Imaging": Supporting information.pdf

**Affiliations**

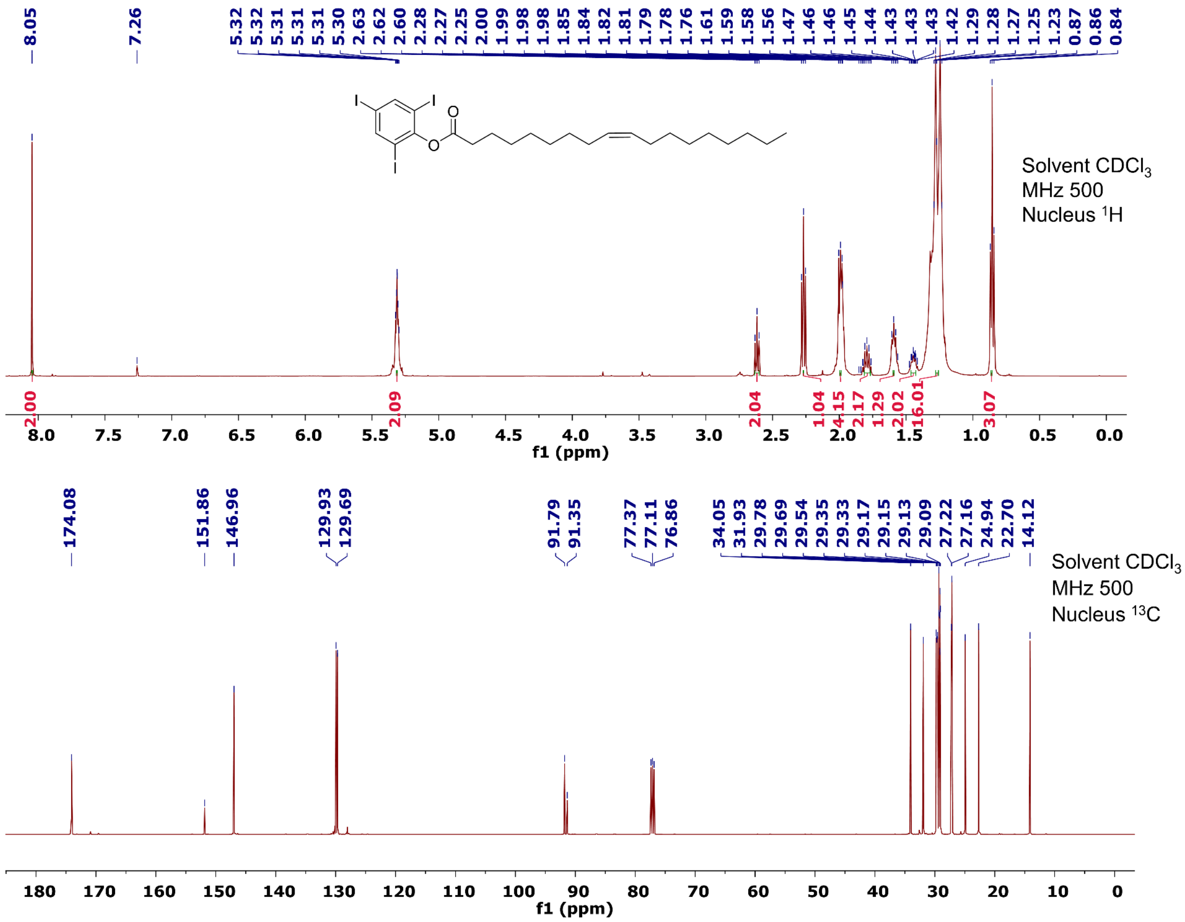


**Supplementary Fig. 1** $^1$HNMR and $^{13}$CNMR analyses of the synthesized TIPhO.

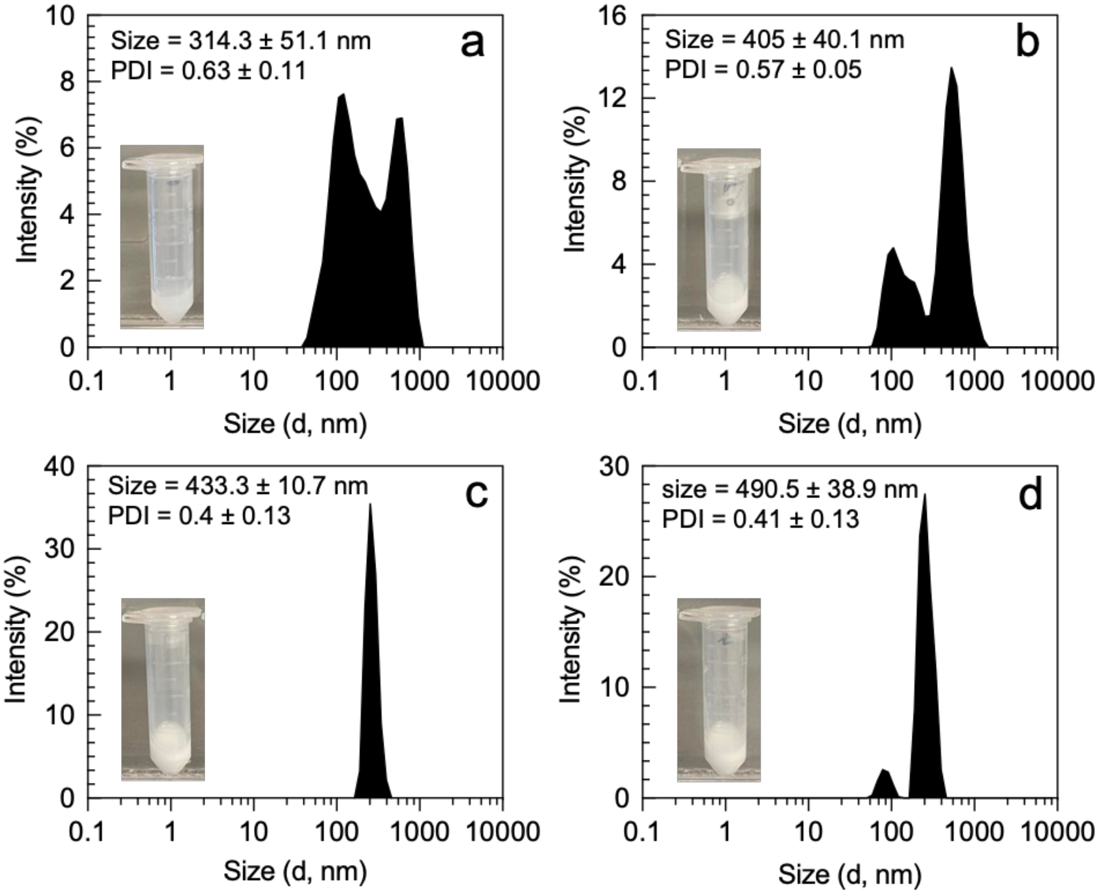


**Supplementary Fig. 2** DLS size histograms of noniodinated LNE1–4 formulations using unmodified oleic acid as a lipid core at different SOR ratios. (a) LNE1 at SOR50, (b) LNE2 at SOR40, (c) LNE3 at SOR30, and (d) LNE4 at SOR20. Larger sizes and PDIs and possible aggregation were obtained which is typically conversely to the TIPhO LNE formulations.

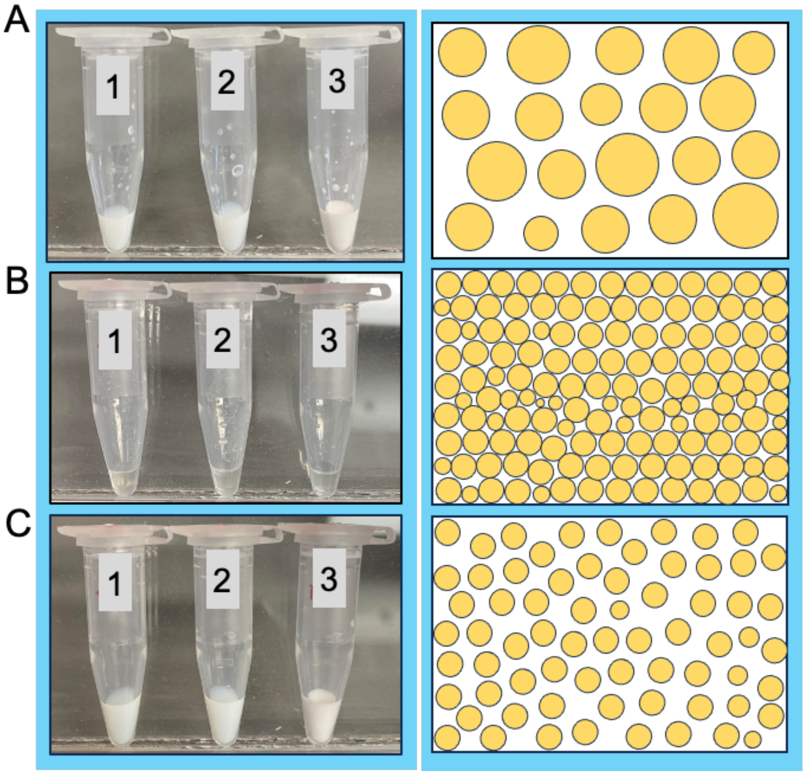


**Supplementary Fig. 3** The appearance of ILNE formulations: (1) ILNE3 with CrEL as a surfactant at SOR30, (2) ILNE3.2 prepared with HS15 CrEL:solutol surfactants (50:50) at SOR30, and (3) ILNE4 with CrEL as a surfactant at SOR40. (A) Fresh samples after preparation immediately. (B) Lyophilized samples (oil phase). (C) reconstitution of lyophilized samples with warm 0.9% normal saline. (Right column): Schematic drawing of the nanodroplet emulsion in suspension in the three phases. The sizes and polydispersity indices (PDIs) of all samples were reduced after the freeze-drying process (see data in **Figure 2d,e** in the main manuscript).

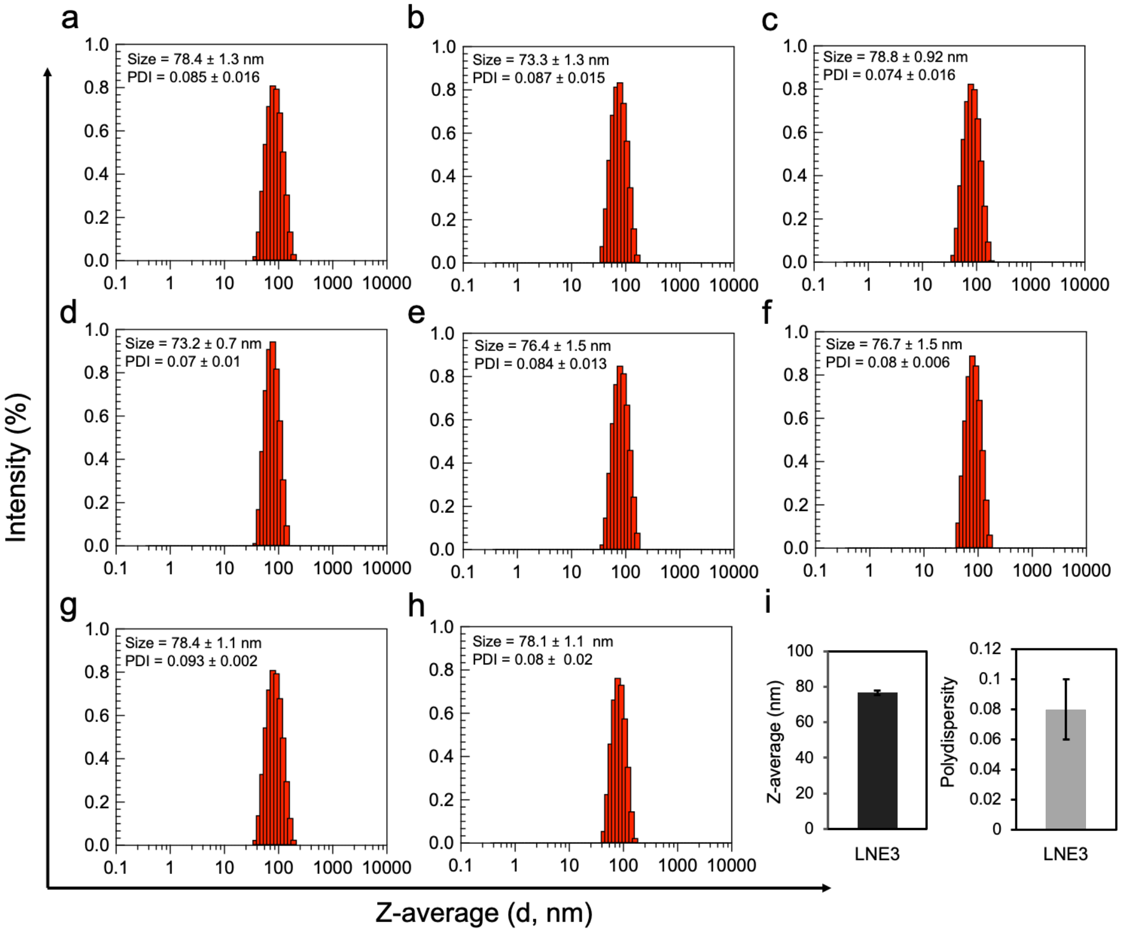


**Supplementary Fig. 4** Batch-to-batch variation of ILNE3: (a-g) DLS histograms of different separate batches of ILNE3 formulations. (h) 35 mL batch of ILNE3. (i). The average size and polydispersity of all batches of ILNE3.

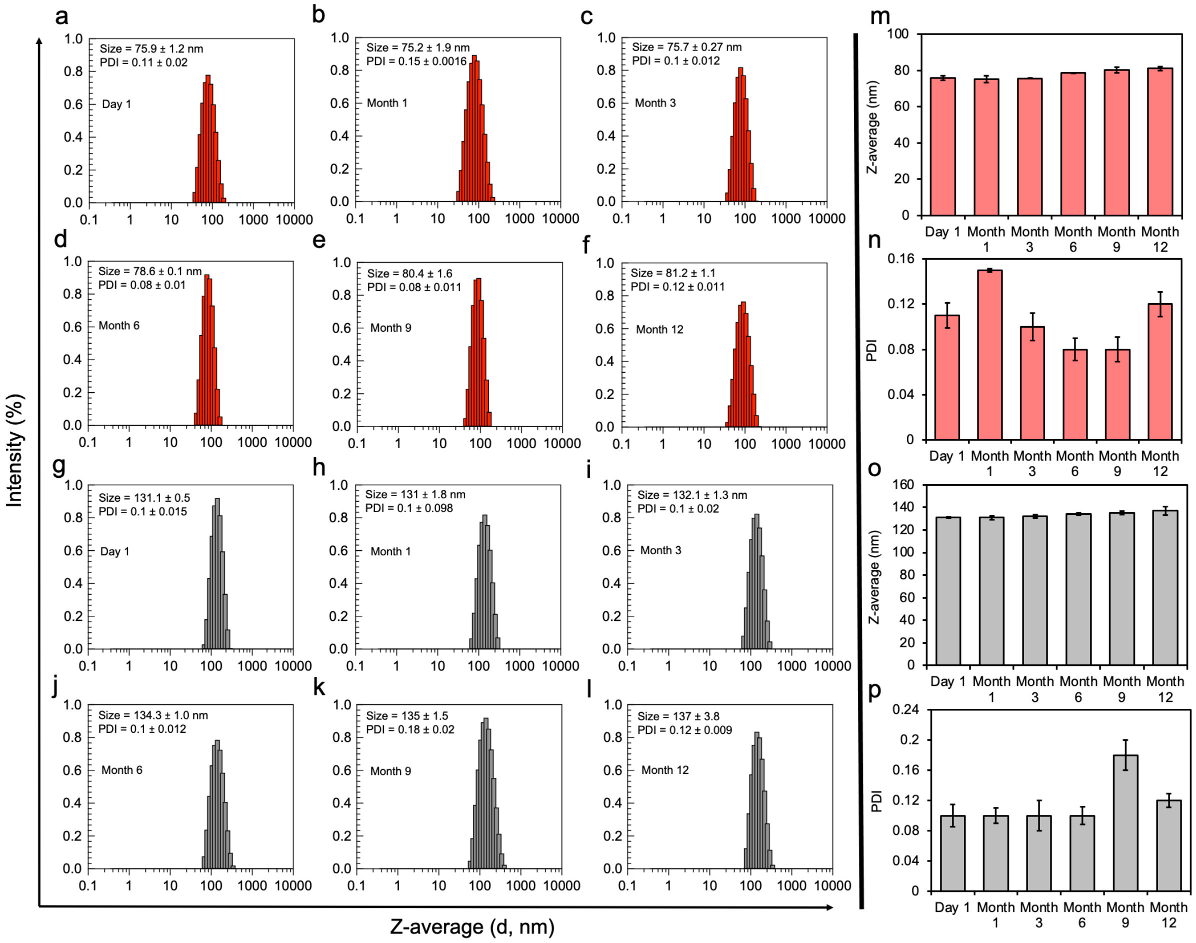


**Supplementary Fig. 5** Shelf-life stability of both ILNE3 (a–f) and ILNE4 (g–l) formulations at room temperature over one-year post-formulation. Average size and PDI of ILNE3 (m and n, respectively). Average size and PDI of ILNE4 (o and p, respectively).

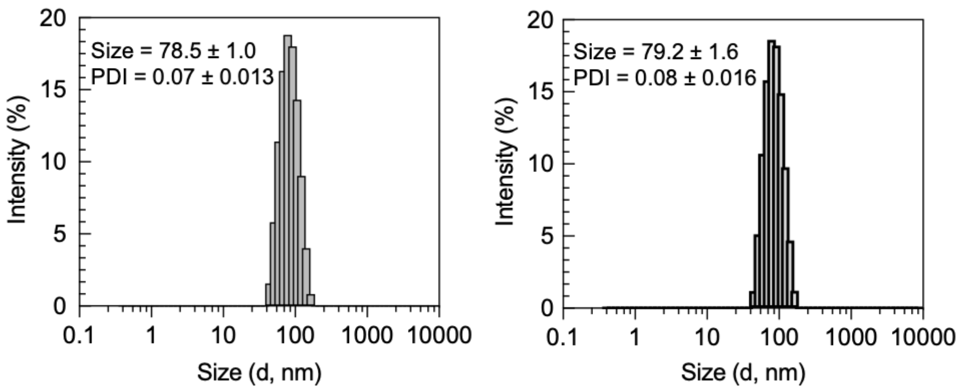


**Supplementary Fig. 6** DLS histograms of ILNE3 stored at 4 °C. (left) Six months post-formulation and (right) nine months post-formulation.

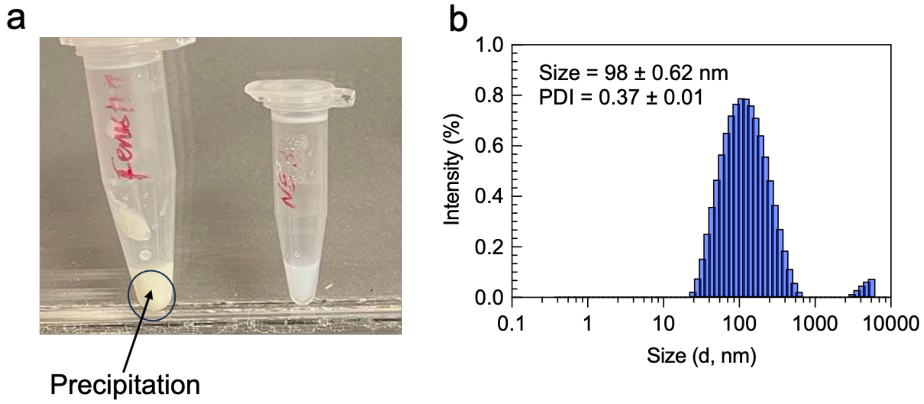


**Supplementary Fig. 7 (a)** Digital photos show the appearance of both the commercial preclinical Fenestra$^{TM}$ HDVC with aggregates and high viscosity (left) and our ILNE3 suspension (right). (b) DLS histogram size of Fenestra HDVC with relatively high PDI.

**Supplementary Table 1** (a) ILNE formulations using PEGylated non-ionic surfactant Solutol® HS15 with TIPhO compound at varying SOR ratios and SOWR40. (b) ILNE formulations using mixed surfactants HS15 and CrEL at a 50:50 ratio, also with TIPhO, at different SOR ratios and SOWR40.

a

| LNE CT agents | SOR | Size (nm) | PDI | Iodinated compound (mg) | Surfactant (mg) | Saline (uL) | Iodine concentration (mg I/mL) Calculated | Iodine concentration (mg I/mL) Measured | 2 mL dose/kg (mgI/kg) | Surfactant (wt/v)% |
|---|---|---|---|---|---|---|---|---|---|---|
| **LNE2.1** | 40 | 92.1±0.96 | 0.22±0.01 | 240 | 160 | 600 | 119 | 140.6 | 281.2 | **16** |
| **LNE3.1** | 30 | 121.8±1.14 | 0.15±0.02 | 280 | 120 | 600 | 139 | 152.5 | 305 | **12** |
| **LNE4.1** | 20 | 152.3±3.3 | 0.2±0.01 | 320 | 80 | 600 | 159 | 176.5 | 353 | **8** |

b

| LNE CT agents | SOR | Size (nm) | PDI | Iodinated compound (mg) | Surfactant (mg) | Saline (uL) | Iodine concentration (mg I/mL) Calculated | Iodine concentration (mg I/mL) Measured | 2 mL dose/kg (mgI/kg) | Surfactant (wt/v)% |
|---|---|---|---|---|---|---|---|---|---|---|
| **LNE2.2** | 40 | 256.7±25.71 | 0.54±0.09 | 240 | 160 | 600 | 119 | 140.6 | 281.2 | **16** |
| **LNE3.2** | 30 | 79.4±0.24 | 0.08±0.01 | 280 | 120 | 600 | 139 | 152.5 | 305 | **12** |
| **LNE4.2** | 20 | 155.7±0.7 | 0.17±0.02 | 320 | 80 | 600 | 159 | 176.5 | 353 | **8** |

**Supplementary Table 2** provides a comparison of the concentrations of non-ionic surfactant CrEL in our ILNE3 formulation versus paclitaxel at two distinct clinical doses (30 and 60 mg/kg of paclitaxel). The comparisons are calculated based on a single dose for each drug. It's worth noting that Taxol (CrEL-paclitaxel) is typically administered in four doses for cancer patients, while our diagnostic drug requires only a single injection. When considering a 4-dose treatment regimen of paclitaxel, the concentration of CrEL in our ILNE3 decreases by **45.6-** and **91.2-fold** compared to that in paclitaxel at the two concentrations (30 and 60 mg/kg), respectively.

| CrEL conc (mg/kg) in ILNE3 | CrEL conc (mg/kg) in Taxol at PTX dose of 30 mg/kg | Reduced amount of CrEL in ILNE3 compared to that in Taxol | CrEL conc (mg/kg) in Taxol at PTX dose of 60 mg/kg | Reduced amount of CrEL in ILNE3 compared to that in Taxol |
|---|---|---|---|---|
| **240** | **2740** | **2740/240 = 11.4 x** | **5480** | **5480/240 = 22.8 x** |

**Supplementary Table 3** Injected volume doses of ILNE3 at three levels of body weight.

| | Per 20g mouse | Per 1kg | Per 70kg patient |
|---|---|---|---|
| **ILNEs** | 40 µL | 2 mL | 140 mL |
| **NP excipients** | 16 mg | 800 mg | 56 g |
| **Iodine (calculated)** | 5.8 mg | 290 mg | 20.28 g |
| **Iodine (found)** | 6.12 | 306 mg | 21.42 g |

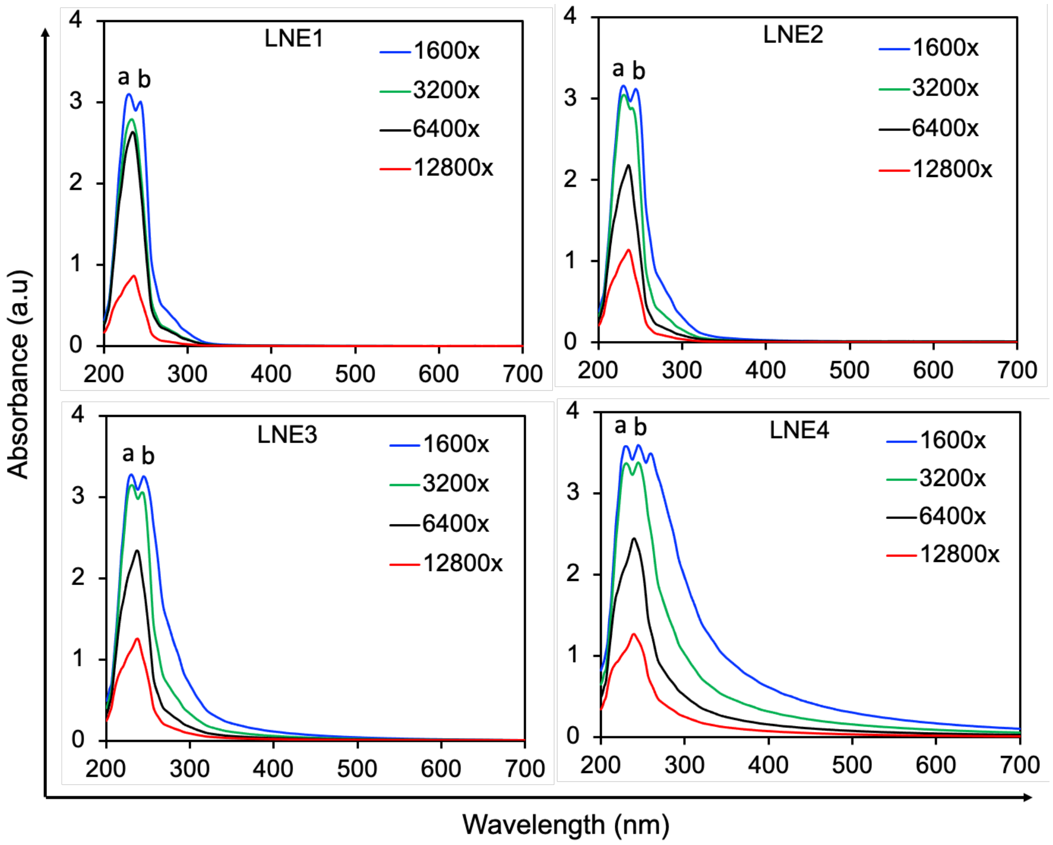


**Supplementary Fig. 8** *UV-vis* absorption spectra of ILNE1–4 formulations at different dilutions, specifically 1600x, 3200x, 6400x, and 12800x). Two distinct peaks were observed for $\lambda_{max}$ at 230 nm (a) for CrEL and 244 nm (b) for TIPhO.

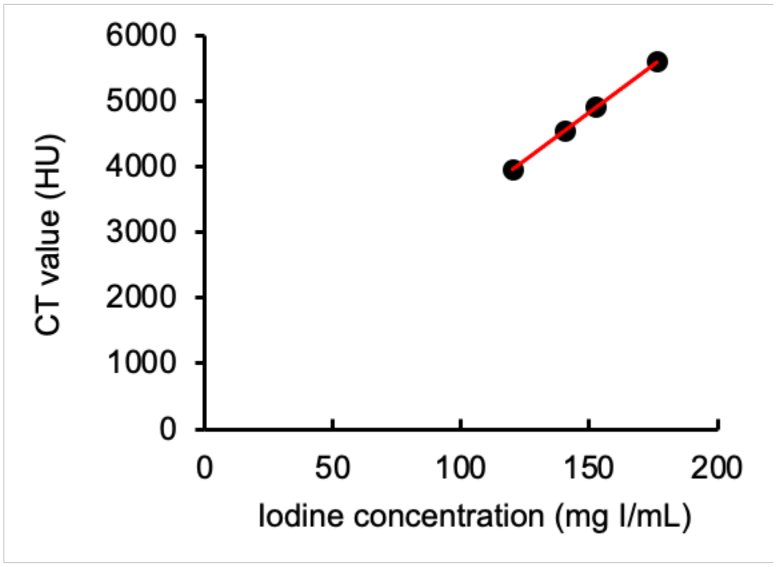


**Supplementary Fig. 9** CT values (HU) of ILNE1–4, with a linear increase of the iodine concentration in the final suspensions.

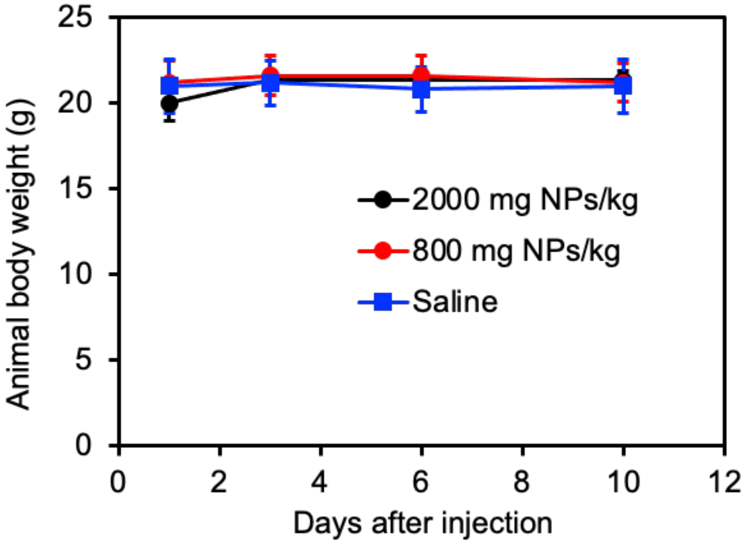


**Supplementary Fig. 10** Animal body weight over time after injection of mice with saline (n =3), ILNE3 at a dose of 800 mg NPs/kg (n=5), and a dose of 2000 mg NPs/kg (n=3).
